# Live Cell Painting: image-based profiling in live cells using Acridine Orange

**DOI:** 10.1101/2024.08.28.610144

**Authors:** Fernanda Garcia-Fossa, Thaís Moraes-Lacerda, Mariana Rodrigues-da-Silva, Barbara Diaz-Rohrer, Shantanu Singh, Anne E. Carpenter, Beth A. Cimini, Marcelo Bispo de Jesus

## Abstract

Image-based profiling has been used to analyze cell health, drug mechanism of action, CRISPR-edited cells, and overall cytotoxicity. Cell Painting is a broadly used image-based assay that uses morphological features to capture how cells respond to treatments. However, this method requires cell fixation for staining, which prevents examining live cells. To address this limitation, here we present Live Cell Painting (LCP), a high-content method based on Acridine orange, a metachromatic dye that labels different organelles and cellular structures. We began by showing that LCP can be applied to follow acidic vesicle redistribution of cells exposed to acidic vesicles inhibitors. Next, we show that LCP can identify subtle changes in cells exposed to silver nanoparticles that are not detected by techniques such as MTT assay. In drug treatments, LCP was helpful in assessing the dose-response relationship and creating profiles that allow clustering of drugs that cause liver injury. Here, we present an affordable and easy-to-use image-based assay capable of assessing overall cell health and showing promise for use in various applications such as assessing drugs and nanoparticles. We envisage the use of Live Cell Painting as an initial screening of overall cell health while providing insights into new biological questions.

## Introduction

Image-based methods for high-content analysis play a pivotal role in uncovering novel mechanisms of action and providing insights into drug-induced phenotypes, offering an overview of cell function (Way et al., 2023). Widely applicable across various biological areas, this approach accelerates drug discovery by offering lots of quantitative and qualitative data, facilitating the identification and optimization of lead compounds (Lin et al., 2020; Zanella et al., 2010), and providing chemical toxicological profiles (Li and Xia, 2019). Image-based profiling assays showcase their versatility by not only predicting cell health outcomes (Way et al., 2021) but also revealing intriguing impacts of genetic perturbations, such as those found by CRISPR-Cas9 editing (Lazar et al., 2023). Additionally, this approach can also identify distinctive epigenetic changes induced by compounds, aiding accurate drug type prediction and even pinpointing drugs associated with reduced tumor regrowth (Farhy et al., 2019). These examples underscore the potential of image-based methods combined with high-content analysis to significantly improve our understanding of biology and to advance applications in various fields, particularly in drug discovery and toxicology.

Furthermore, image-based methods offer a non-invasive means of studying cellular dynamics, allowing for real-time observations and analysis of dynamic processes within cells. Temporal evaluation improves the ability to capture transient events, enabling a more comprehensive understanding of cellular responses (André et al., 2023; Garvey et al., 2016). Applying machine learning algorithms to image analysis enhances the accuracy of extracting meaningful information from complex datasets, further advancing the potential of image-based methods (Siegismund et al., 2022). Finally, the combination of image-based profiling with multi-omics data, such as genomics and transcriptomics, provides a holistic understanding of cellular responses, linking morphological changes to underlying genetic and molecular alterations (Watson et al., 2022). These examples showcase the versatility of image-based methods, which can provide an invaluable tool for studying dynamic cellular processes.

Image-based assays on fixed cells are straightforward and convenient for staining and imaging a large number of samples, but this approach has some limitations. The physical processes of cell fixation and permeabilization can lead to image artifacts, significantly altering intracellular structures, including protein distribution, and causing the loss of chemical gradients (Schnell et al., 2012). While fixed-cell imaging involves the preservation of cellular structures at a specific moment in time, live-cell image-based methods allow for detailed temporal examination of cellular components, such as organelle activity (*e.g.*, vesicles acidification, nucleoli organization) and cytoskeletal (Selase, 2020). In addition to real-time observation, live-cell imaging facilitates the monitoring of cellular responses to external stimuli over extended periods, providing insights into the kinetics and dynamics of cellular processes (Gordonov et al., 2016). This is particularly valuable in understanding long-term effects and cellular responses, such as those occurring in chronic diseases or during prolonged drug exposure (Tolosa et al., 2019).

Advances have been made in the field of live-cell imaging through the development of new technologies, but some limitations still persist. Establishing cell lines that express fluorescent reporter proteins often requires the constant use of antibiotics, which can result in artifacts by perturbing cell metabolism (Binó et al., 2022; Cuny et al., 2022). Alternatively, using gene-editing technologies offers more precise methods for introducing fluorescent reporters without the need for antibiotics, thereby addressing some challenges associated with live cell imaging. However, the widespread use of this technology still faced some limitations, such as large array sizes, the lack of an efficient gene delivery system and high associated costs (Huang et al., 2023). Although some approaches enable live cell profiling, there remains an urgent need for new assays that are both user-friendly and cost-effective.

Here, we present Live Cell Painting, which uses acridine orange (AO) to stain live cells for image-based profiling. Acridine orange metachromatic fluorescence can report on different cellular chemical microenvironments. AO monomers intercalate between neighboring pairs of DNA and RNA, absorbing blue light (∼458 nm) and emitting green light (∼530 nm). At higher concentrations in cells, AO interacts electrostatically with DNA and RNA and can also organize into stacks and accumulate in acidic vesicles. When AO is organized as stacks it will shift the emission spectra to red light (∼640 nm). To establish Live Cell Painting, we optimized AO concentration, evaluated its versatility across different cell lines, and determined non-toxic concentrations. We next validated acridine orange affinity for acidic vesicles and the ability to track vesicle redistribution and cell phenotype assessment. To demonstrate image-based profiling capacity and versatility, we used different perturbations: silver nanoparticles and drugs. Using silver nanoparticles, we demonstrated the ability of Live Cell Painting to assess the dose-response relationship. For drug assessment, cells were treated with selected compounds from the Drug-Induced Liver Injury (DILI) rank list, and data were grouped into profiles based on images depending on concentration and compound. Our results demonstrate that Live Cell Painting holds promise as an easy-to-use method for image-based profiling and analysis of individual features in living cells, providing valuable insights into cell health and condition.

## Results

### Live Cell Painting: broadly applicable to cell lines and cytotoxicity assessment

To establish the Live Cell Painting for image-based profiling, we began by assessing the cytotoxic profile of cells exposed to AO using the CellTiter-Glo^®^ Luminescent Cell Viability Assay. Our goal was to identify an optimal AO concentration that ensures cell viability while allowing effective staining of nucleic acids and acidic vesicles. For this purpose, we selected four cell lines: Huh-7 (hepatocarcinoma), MCF-7 (breast cancer), PNT1A (non-cancer prostate epithelium), and PC-3 (prostate cancer) (Supplementary Figure S1). Cells were treated with increasing concentrations of AO (2.5, 5.0, 10.0, 20.0, and 40.0 μM) for 4 hours, estimating the time cells would be exposed to AO during image acquisition.

Cells exhibited distinct tolerances when exposed to AO. Huh-7 cells tolerated up to 20 μM of AO for 4 hours, showing no significant difference (p > 0.05) compared to the negative control (Figure 1A, data normalized to untreated cells). At higher concentrations (40 μM, corresponding to 1.6 on the logarithmic scale, Figure 1A), cell viability was significantly reduced (p < 0.05) to approximately 40%. Therefore, we chose 20 μM to proceed with the following experiments. MCF-7 cells tolerated higher concentrations of AO, as cell viability was only significantly reduced (p < 0.05) to approximately 70% at 40 μM AO, revealing a cell type-dependent response (Figure 1B). Prostate cells (PNT1A and PC-3) showed even lower tolerance to AO exposure; the maximum AO concentration that did not affect cell viability was 2.5 μM (Supplemental Figure S1). In Figure 1B, the AO-Green channel allows us to visualize DNA and RNA, allowing for identification of the nuclei and cell objects, while the AO-Red channel highlights acidic vesicles. This dual-channel approach provides insights into the cellular structures and functions under investigation. We conclude that cell tolerance to AO is cell type dependent; therefore, it is imperative to evaluate cell viability to determine the AO concentration with minimal toxicity while maintaining its ability to stain cell compartments.

**Figure 1.**
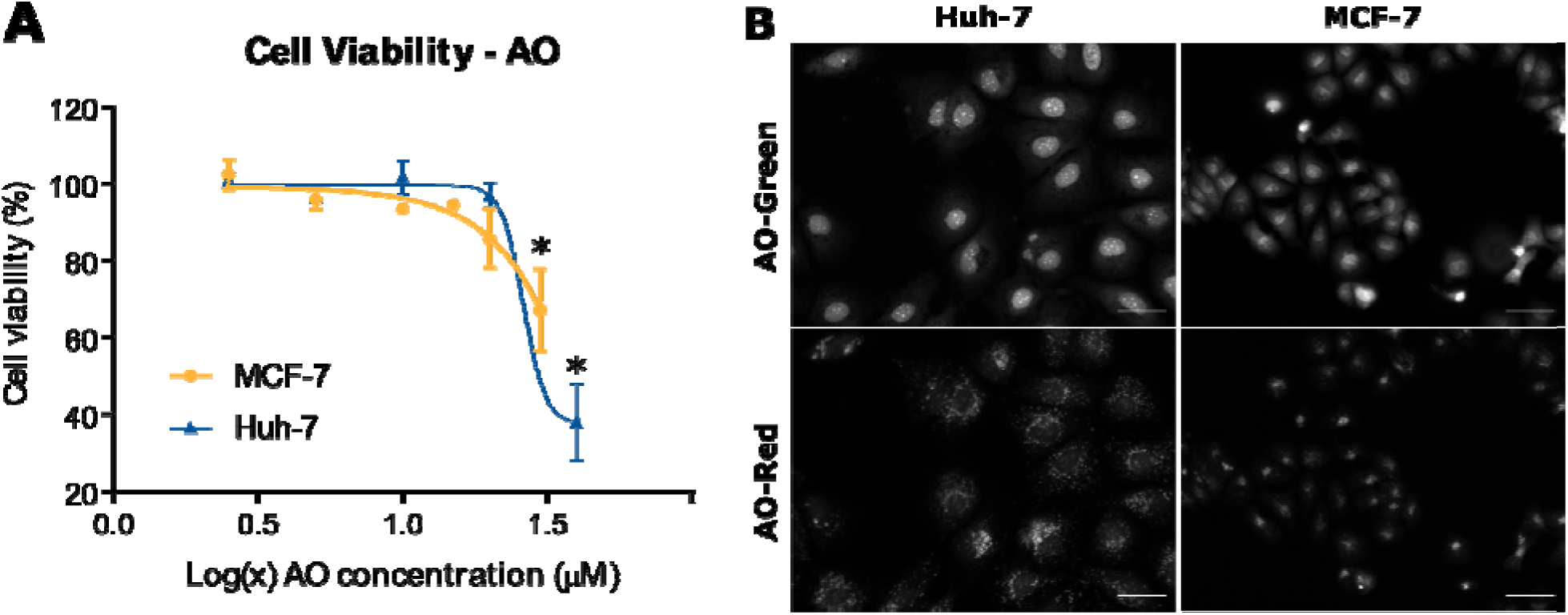
Cell viability in the presence of AO assessed by ATP concentration. Huh-7 and MCF-7 cells were plated at a density of 8000 cells/well in a 96-well plate. After 24 hours, cells were treated with AO (2.5, 5.0, 10.0, 20.0, and 40.0 μM) for 4 hours. Then, viability was assessed using CellTiter-Glo^®^ by measuring ATP concentration with luminescence. ATP concentration in the cells was determined by fitting a linear function to the ATP standard curve and calculating the relative viability compared to the negative control. (A) Viability curve for Huh-7 and MCF-7 cells exposed to AO and (B) Representative images were acquired (Cytation 5, 20x objective) for Huh-7 and MCF-7 cells exposed to 20 and 10 μM of AO, respectively, represented by the AO-Green and AO-Red channels. Scale bar = 50 μm. Statistical test performed with two-way ANOVA followed by Sidak’s multiple comparisons test: * p < 0.05.

### Acridine Orange captures the dynamics of acidic vesicles

To validate AO accumulation in acidic vesicles (late endosomes and lysosomes) and to illustrate that AO can be used to detect vesicle (re)distribution, we used two well-known acidification inhibitors: Bafilomycin A1 and Chloroquine. Our underlying hypothesis was that treatment with these inhibitors would result in a reduction in fluorescence intensity and potentially alter the distribution of vesicles in the cytoplasm. To assess this hypothesis, we incubated Huh-7 cells with inhibitors (Bafilomycin A1 1 µM and Chloroquine 50 µM) for 0.5 h or 4 h and used either Lysotracker Deep Red or Acridine Orange to follow acidic vesicle distribution. Lysotracker wa used to confirm the presence of acidic vesicles in control cells and validate the decrease in fluorescence intensity upon inhibitor treatment (Supplemental Figure S2).

Vesicle morphology changed upon incubation with the inhibitors, showing changes in the distribution, number, and intensity of acidic vesicles over time (Figure 2B). After segmentation of cell borders, the total intensity of AO across all cells was calculated by multiplying the cell number by the mean AO-Red cell intensity (Figure 2A). Bafilomycin A significantly decreased overall intensity at 0.5 h and 4 h (p < 0.05). Intriguingly, Chloroquine significantly increased the total intensity after 0.5 h of treatment and subsequently decreased to levels similar to those of untreated cells after 4 h of treatment. AO begins to accumulate in the cell nuclei after 4 h of treatment with Bafilomycin A and Chloroquine (Figure 2B). This profile differs from the untreated cells, indicating a subtle toxic effect after 4 h.

**Figure 2:**
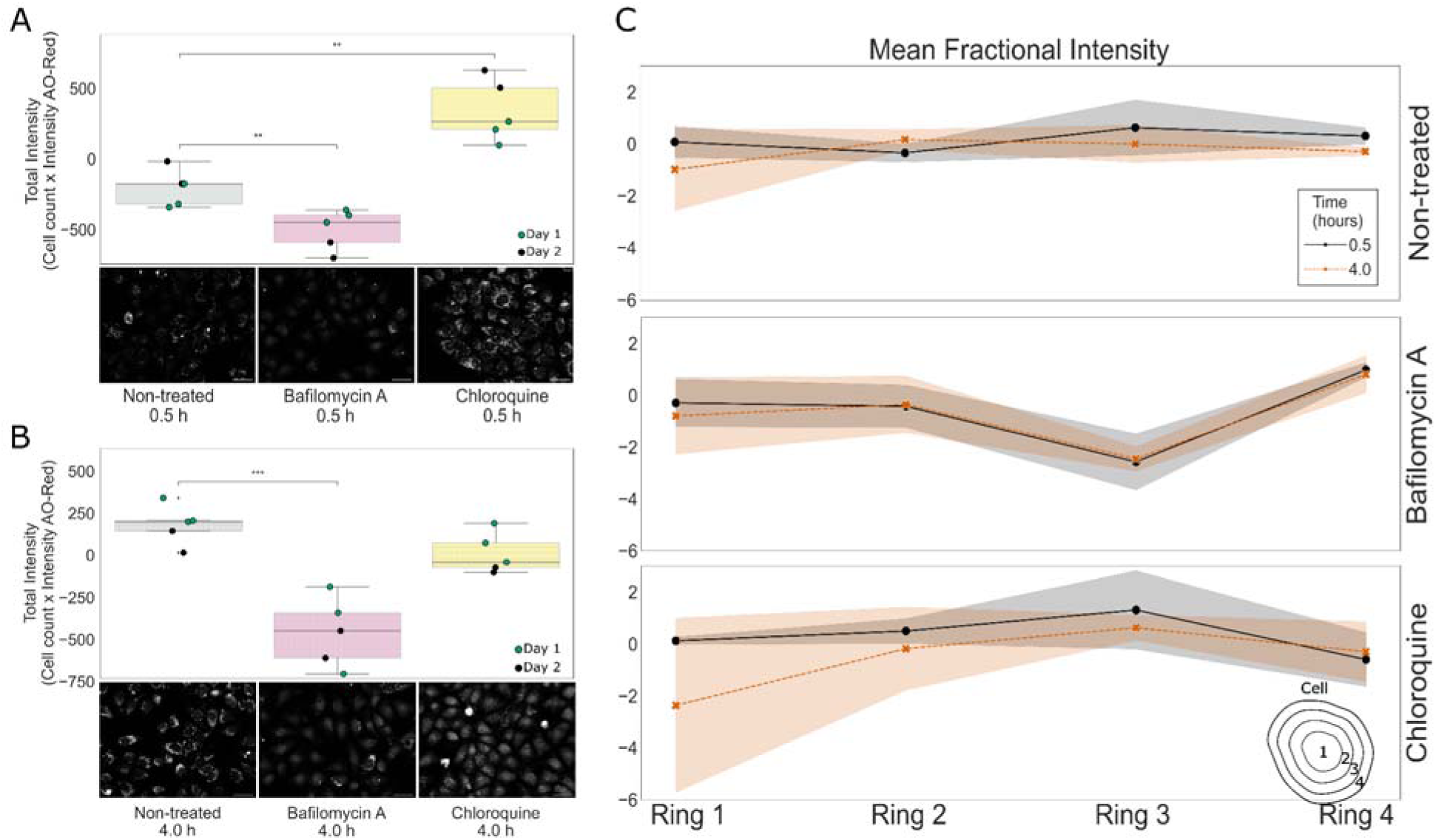
Acridine orange assessment of acidic vesicle dynamics in Huh-7 cells. Huh-7 cells were plated at 8000 cell per well into 96-well plates. The next day, cells were treated with Bafilomycin A 1.5 μM + AO or Chloroquine 50 μM + AO for 0.5 hours or 4 hours. Subsequently, the medium was replaced with Fluorobrite, and images were captured using the Cytation 5 Biotek system, with a 20x objective. (A-B) The total intensity was calculated by the multiplication of cell count and the cell intensity in the AO-Red channel, where vesicles are identified. Each point represents data aggregated per well, colored by batch. Below each graph are representative images of Huh-7 cells in the presence and absence of inhibitors at the respective exposure time (0.5 or 4 hours). Scale bar = 50 μm. (C) Each identified cell was split into 4 concentric rings using CellProfiler, and the intensity was measured in each concentric ring. The mean fractional intensity is represented in each concentric ring (1 of 4, 2 of 4, 3 of 4, and 4 of 4). The intensity at each concentric ring is shown in the x-axis, starting from the center of the cell (1 of 4) to the border of the cell (4 of 4) Experiments were performed in two independent replicates, n = 3 wells. Statistical test (independent t-test) was performed using the statannotations library. * p statistical difference < 0.05.

The Live Cell Painting assay can also report the patterns of acidic vesicle distribution across the cell. To assess acidic vesicle distribution across the cell, we used a CellProfiler module called MeasureObjectIntensityDistribution, which divides cells into four concentric ring and measures the fraction of the cell’s staining intensity in each ring in the AO-Red channel (Figure 2B). For Bafilomycin A1, the intensity at ring 3 decreased at 0.5 and 4 hours, revealing a pattern of change in the morphology of the treated cells. Chloroquine shows a similar pattern of vesicle distribution compared to untreated cells, while Bafilomycin A1 shows a distinct distribution that persists throughout 4 h of treatment (Figure 2C). Collectively, these finding suggest that AO successfully captured the cellular state of acidic vesicles and this approach can provide fine details about cell response to perturbation.

### Acridine Orange captures dose-response effects

To illustrate how AO can be used to identify perturbations to cells caused by dose-response treatments, we treated cells with increasing concentrations of silver nanoparticles for 24 h. Nanoparticles exploit unique properties at the nanometer scale and provide solutions to various problems, particularly in areas such as nanomedicine. However, their use also poses challenges due to possible undesirable effects, in particular toxic effects when reaching eukaryotic cells. Here, we used nanoparticles in two sizes: 40 nm (referred to as Ag40, Supplementary Figure S3) and 100 nm (referred to as Ag100). The t-SNE algorithm was used for dimensionality reduction and visualization (Figure 3A), providing insight and intuition into different AgNP treatment patterns. Higher concentrations of Ag100 are located at the top left, while lower concentrations are located at the bottom right, suggesting that AO has the potential to identify dose-response changes. We also compared the sensitivity of Live Cell Painting with the MTT assay, a traditional method for assessing cell viability. Concentrations detectable by Live Cell Painting were up to 40 times lower than the IC_10_, based on values previously determined with MTT (*i.e.*, the inhibitory concentration of 10% of the population), indicating a greater sensitivity.

**Figure 3.**
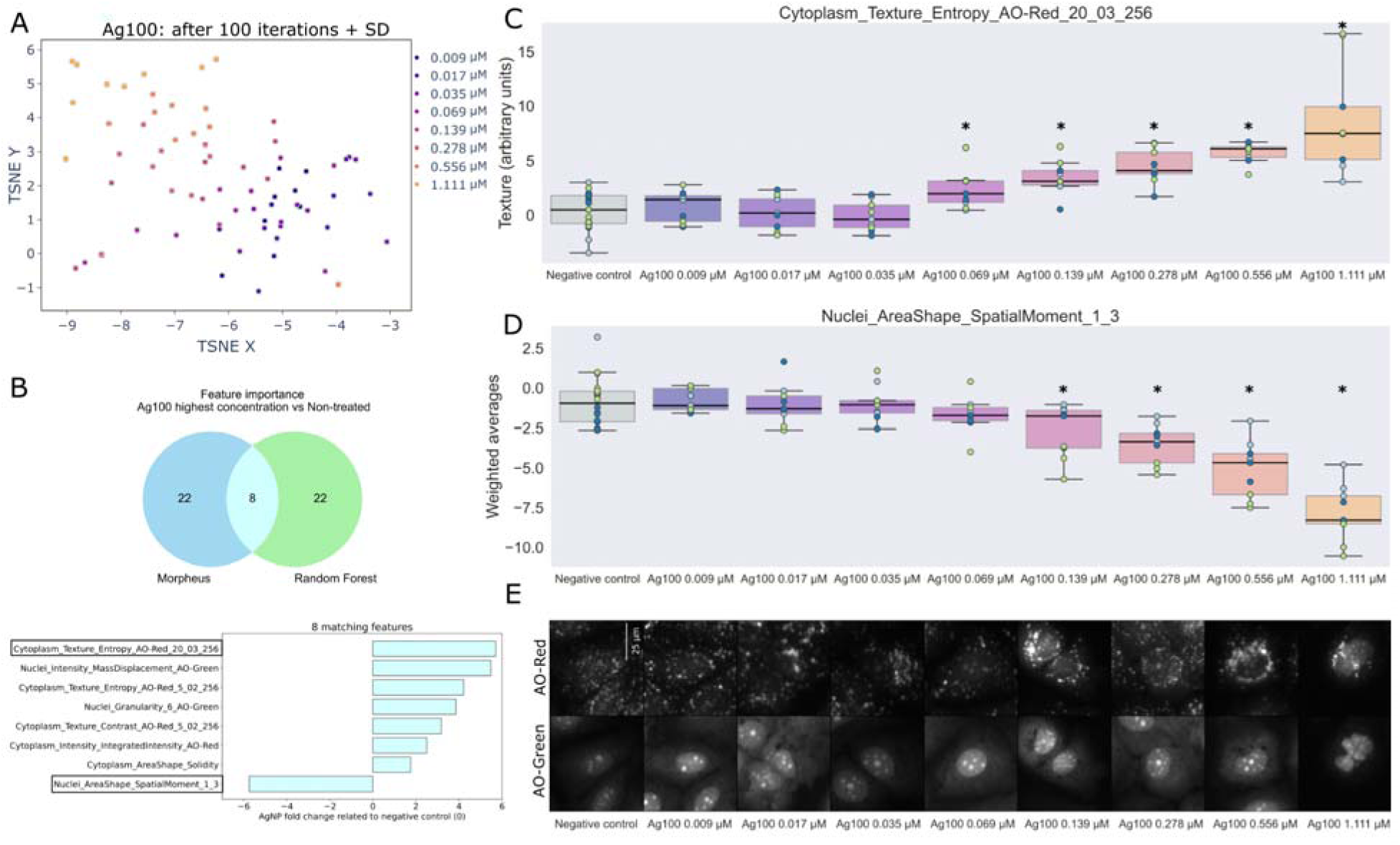
Acridine Orange captures the dose-response effect induced by silver nanoparticle (AgNPs) on Huh-7 cells. Huh-7 cells were treated with Ag100 (nanoparticle size 100 nm) at increasing concentrations for 24 h. Then the images were acquired after AO staining (Cytation 5, 20x objective). Objects were segmented using Cellpose software and features were extracted using CellProfiler software. Next, features were aggregated by well, normalized, and selected using Pycytominer and batch corrected using pyComBat. (A) t-SNE was used for dimensionality reduction and visualization. After 100 iterations, the mean and standard deviation embedding were calculated. (B) Random forest ran for 100 iterations to differentiate between two classes: negative control versus combining Ag100 0.556 and 1.111 μM. Then, the mean feature importance and standard deviation were calculated. Morpheus software was used to perform marker selection between the same random forest classified groups, and the top 30 features were selected and compared. Among those, 8 features were consistent between methods and were plotted to show the fold change compared to the negative control. The two features highlighted by black boxes are presented in C and D; (C) The intensity of AO-Red in the nuclei; (D) The spatial moment of the nuclei, representing the shape, size, rotation, and location of the nuclear object, and (E) Representative images were retrieved from the dataset using k-means clustering (Garcia-Fossa et al., 2023). Experiments were performed in three independent replicates, n = 3 and each point in the plots consists of well-aggregated data, and the points are colored by the batch in C and D. Independent t-test calculated using the statannotations library. * p statistical difference < 0.05. Scale bar = 25 μm.

To confirm the dose-response profiles observed in the t-SNE representation, we used two approaches to identify the most important features that distinguish between negative control and the highest concentrations of Ag100. First, we used Morpheus software’s (https://software.broadinstitute.org/morpheus/) marker selection tool to perform a t-test and provide the 30 most important features. Additionally, we used Random Forest to rank features by their importance using bootstrapping. Among the 30 most important features, eight features matched between the models. For these features, we plot their fold change relative to the negative control (Figure 3B), ranging up to 6 times higher or lower than the negative control. To further investigate the dose-response effect on specific features, we chose the two features that had the highest negative or positive response related to the negative control (the first and last feature in the y-axis in Figure 3B), Cytoplasm_Texture_Entropy_CorrPI_20_03_256 and Nuclei_AreaShape_SpatialMoment_1_3 (Figure 3C and D) and identified a strong dose-response effect to increasing Ag100 concentrations. The six remaining features which also presented strong dose-response behavior are available in Supplemental Figure S4.

The representative images (Figure 3E), retrieved using k-means clustering (Garcia-Fossa et al., 2023), illustrate substantial effects on cell size and intensity in both channels, in cells treated with Ag100. From concentrations of 0.139 μM onward, the size of the cells begins to shrink, which is reflected in the irregular shape of the nuclei. This can be quantified using the SpatialMoment function, which is a series of weighted averages that provide information about the object’s shape, dimensions, orientation, and position (Figure 3D). Interestingly, both features reveal statistically significant changes in the cells at lower dose points than the IC_50_ previously determined by the MTT assay (Ag100 IC_50_ determined by MTT was 0.042 μM versus 2.937 μM determined by fitting a curve to Figure 3C). These results demonstrate that the Live Cell Painting successfully provided dose-response analysis and, based on morphological changes, exhibited greater sensitivity than the MTT assay.

### Live Cell Painting for profiling hepatocytes undergoing Drug-Induced Liver Injury

After showing that Live Cell Painting captures details about intracellular organization and that this could be sensitive enough to reveal how cells respond to perturbations, we hypothesized that Live Cell Painting could be used in image-based profiling. To test this hypothesis, we treated Huh-7 cells with six compounds present in the Drug-Induced Liver Injury (DILI-Rank) dataset (Research, 2020). We then tested whether the cells could be grouped by the compound based on the induced phenotype in hepatocytes. The drugs (1,036) are divided into four classes based on their potential to cause liver injury: no DILI concern, less DILI concern, major DILI concern, and equivocal DILI concern. We used lactose and untreated cells as negative controls. Among DILI compounds, six were selected and used to treat cells for 24 h (Table 1); cells were then stained with AO and prepared for imaging.

**Table 1.** Compounds used to treat cells for the DILI drug screening assay, concentrations used, and described mechanism of action. The DILI-concern and severity class were based on the DILIrank dataset (Chen et al., 2016).

| Compound | Concentration (μM) | Mechanism of action | DILI-concern/<br>severity class |
| --- | --- | --- | --- |
| Untreated cells | 0 | Negative control | - |
| Lactose | 1 and 10 | Negative control | - |
| Aspirin | 1 and 10 | COX inhibitor (Henry et al., 2017; Yang et al., 2017) | Less-concern/class 0 |
| Orphenadrine citrate | 1 and 10 | histamine/NMDA receptors inhibitor (Kornhuber et al., 1995; Rumore and Schlichting, 1985) | Less-concern/class 0 |
| Etoposide | 1 and 10 | Topoisomerase II inhibition (Montecucco et al., 2015) | Less-concern/class 3 |
| Lovastatin | 1 and 10 | HMG-CoA reductase inhibitor (Duong and Bajaj, 2023) | Less-concern/class 3 |
| Cyclophosphamide | 1 and 10 | DNA alkylation (Ogino and Tadi, 2023) | Less-concern/class 5 |
| Amiodarone | 0.2 and 2 | sodium/potassium-ATPase inhibitor (Anthérieu et al., | Most-concern/class 8 |

All cells treated with the compounds were phenotypically active, meaning that their replicate image-based profiles are statistically retrievable relative to negative controls (mean Average Precision (mAP) > 0.05, except for orphenadrine 1 μM which was filtered out) (Kalinin et al., 2024). Some compounds formed more distinct clusters than others. For example, orphenadrine and etoposide showed clear clustering as detected by Live Cell Painting profiling (Figure 4A), which was further confirmed by unsupervised machine learning algorithm clusters based on density (Figure 4B). The clustering was not determined by the DILI severity class, where only the 0-class is identified as a cluster (Figure 4C). Representative images of single cells treated with DILI drugs and negative control were retrieved from the dataset using k-means clustering (Garcia-Fossa et al., 2023) (Figure 4D).

**Figure 4.**
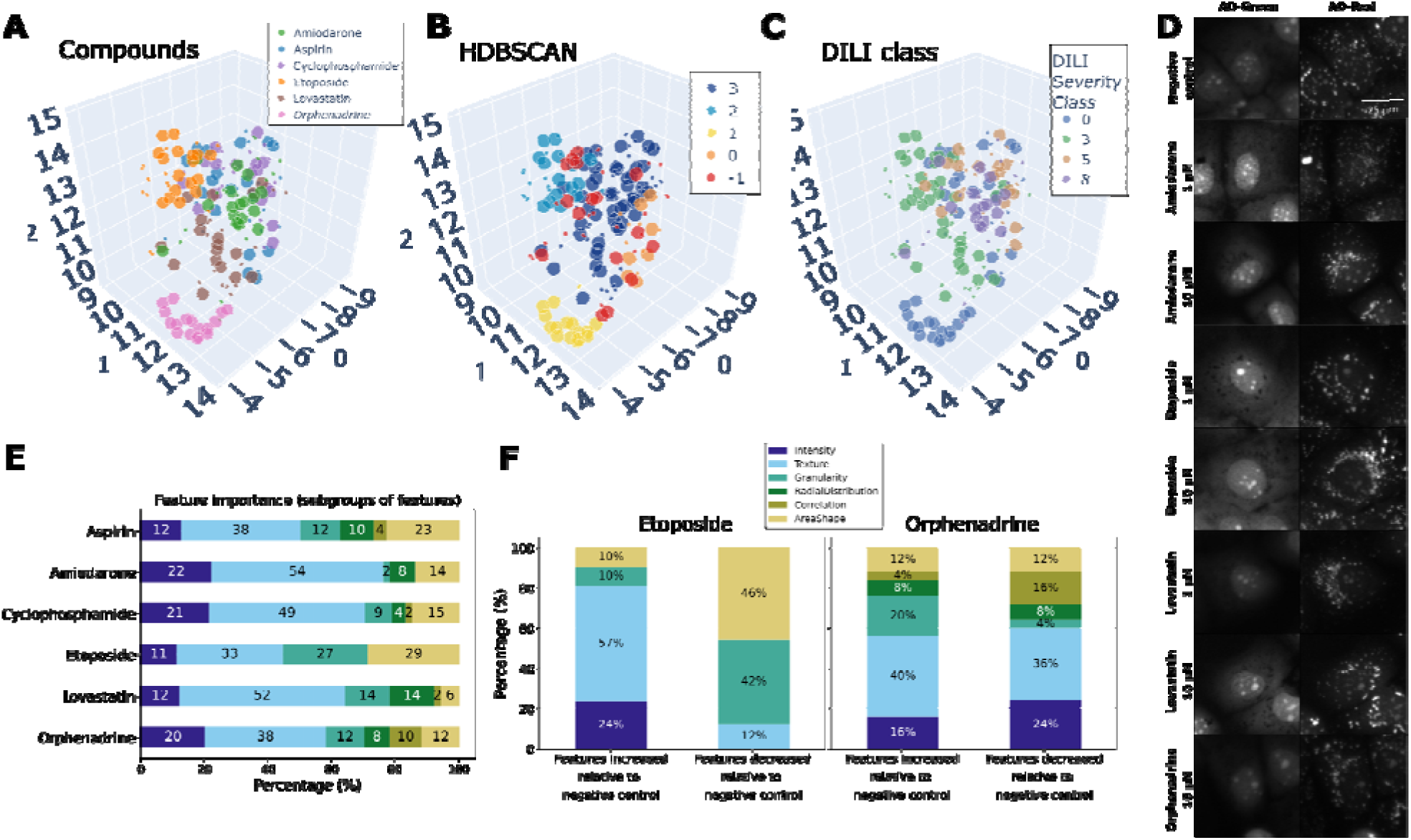
Live Cell Painting captures the phenotypic profile of Huh-7 cells treated with drug from the Drug-Induced Liver Injury (DILI) rank. Huh-7 cells were treated with drugs at the indicated concentrations (Table 1) for 24 hours. After that, cells were stained with AO, and images were acquired using the Cytation 5 20x objective in the AO-Green and AO-Red channels. The images were processed with cell segmentation and feature extraction using Cellpose and CellProfiler. Next, the data were aggregated by well, normalized, and feature selected using Pycytominer and batch corrected using pyCombat. (A) UMAP (cosine metric) was used to reduce the dimensionality of approximately 300 features. The data was color-coded for each drug and the size of the dots represents the dose of the compounds (lower concentrations are smaller, higher concentrations are larger); (B) Samples were classified using unsupervised HDBSCAN clustering (Hierarchical Density-Based Spatial Clustering of Applications with Noise). Noisy samples are given the label -1; (C) UMAP labeled by DILI severity class; (D) Representative images of single cells; (E) Matching features between random forest and Morpheus (50) were categorized into subgroups, and (F) Features increased or decreased relative to the negative control were categorized into subgroups for etoposide and orphenadrine. The experiments were performed in five independent batches with 4 replicates each.

To examine the features captured by Live Cell Painting able to distinguish treated cells from the negative control, we used the same approach as in the section above. We identified the 100 most important features using random forest ensembles and Morpheus t-test for each compound against the negative control, resulting in 50 features matching between the Random Forest and Morpheus approaches. We found that texture-related features were important to determine the profile of the DILI compounds, followed by Intensity, AreShape, and Granularity (Figure 4E). Therefore, we show that Live Cell Painting captures the phenotypic profile of DILI-ranked drugs and provides phenotypic pathway visualizations for each compound.

To understand the patterns of morphological effects in which features were altered by the treatments, we group the features into six categories (AreaShape, Intensity, Texture, Correlation, Granularity, and RadialDistribution), as well as features separated by channel (AO-Green, AO-Red, and AreaShape) (Supplemental Figure S9). Of the top 50 features, 52% belong to the AO-Red channel when cells are treated with amiodarone and 90% in cells treated with lovastatin (Supplemental Figure S9). Despite amiodarone and lovastatin having different mechanisms of action (Table 1), their phenotypic profiles are similar. They are classified into the same class by HDBSCAN, an unsupervised clustering algorithm based on density (Figure 4B, class 3). These two compounds share 30% of their most important features (Supplemental Figure S9), all of which belong to the AO-Red channel and are largely related to texture and radial distribution of intensity in the AO-Red channel, which may be linked to changes in the distribution of acidic vesicles. Another similarity is that amiodarone and lovastatin induce direct or indirect disruption of the mitochondrial respiratory chain (Donato and Tolosa, 2021), leading cells to present a similar phenotype.

For compounds that presented a stronger profile - etoposide and orphenadrine - we split the important features into higher or lower values relative to the negative control, categorizing them as increased or decreased compared to the baseline. The features were then analyzed regarding compartment, channel, feature subgroup, and their fold change related to the negative control (Figure 4F). Both compounds induce different phenotypes that are distinguishable in the representative images (Figure 4D) and also in the clustering, each assigned to a class (Figure 4B, class 1 for orphenadrine and class 2 for etoposide). Etoposide, a topoisomerase II inhibitor, induces significant nuclear structural changes characterized by increased texture correlation and granularity in nuclei (Figure 4F and Supplemental Figure S11). Additionally, treated cells exhibit a fivefold increase in Cytoplasm_AreaShape_Perimeter, indicating cellular enlargement (Supplemental Figure S11D). Orphenadrine binds and inhibits both histamine H1 receptors and NMDA receptors, and it is classified as safe for the liver (Class 0 in DILI severity). Analysis of its effects on Huh-7 cells revealed elevated values in vesicle area shape and eccentricity, suggesting potential alterations in vesicular trafficking or cellular secretory pathways. Conversely, reduced intensity measures in nucleoli and cytoplasmic mass displacement may indicate lowered metabolic activity or changes in protein synthesis (Figure 4F and Supplemental Figure S12). These adaptations likely represent responses to cellular stress or shifts in metabolic demands within liver cells. By linking these features to specific aspects of etoposide’s and orphenadrine’s action, we can better understand the cellular responses and validate the use of AO for assessing the drug’s effects. These results demonstrate that Live Cell Painting can be a useful tool for determining the phenotypic profile in living cells and provides mechanistic insights into visualizing phenotypic pathways for each activity group.

## Discussion

Here we propose the Live Cell Painting assay, an informative image-based assay that can be performed in living cells using Acridine Orange (AO) as a fluorescent dye. Leveraging the dual-emission property of AO allows for distinguishing many intracellular structures, including nucleic acids and acidic compartments such as late endosomes and lysosomes. We show that Live Cell Painting provides a wealth of information by monitoring the morphological changes of cell populations in response to perturbations such as drugs or nanoparticles. With these perturbations, we have gained additional insights into the Live Cell Painting, highlighting its capacity to cover a wide range of phenotypes, such as clustering DILI drugs. Additionally, it was highly sensitive to detect perturbations, such as those at low concentrations of nanoparticles, which are crucial for several processes reflected in cell morphology.

We found that Live Cell Painting can be used to study specific cellular mechanisms such as acidic vesicle dynamics. AO staining captures details of acidic vesicles distribution, demonstrated by using acidification inhibitors and after analyzing the AO-Red intensity within concentric rings in the cell. Bafilomycin A1, which inhibits autophagosome-lysosome fusion and acidification (Yoshimori et al., 1991), significantly reduced the number and intensity of acidic vesicles in Huh-7 cells, consistent with its role in preventing autolysosome formation. Conversely, chloroquine, known for its inhibition of autophagy and lysosomal function (Homewood et al., 1972), induced phospholipidosis in hepatocytes. This condition was characterized by the accumulation of phospholipids in lysosomes, resulting in stronger AO staining of acidic vesicles compared to untreated cells (Cho et al., 2020) These findings demonstrate the assay’s capability to capture specific morphological details of intracellular events induced by pharmacological interventions.

Furthermore, Live Cell Painting revealed dose-response effects in cell responses to silver nanoparticles (AgNP) and other perturbations. This approach uses a single dye as opposed to other assays such as the original Cell Painting assay (Bray et al., 2016; Cimini et al., 2023) which requires six dyes added in two staining steps. Our method has demonstrated its ability to uncover subtle changes in treated cells that are often missed by conventional techniques such as MTT assays (Supplemental Figure S6). In our dose-response assay with low concentrations of AgNP, we observed alterations in acidic vesicle distribution and cell size, potentially linked to autophagy pathways. Previous studies on various cell types and nanoparticles suggest that cells might utilize the endo-lysosomal pathway to process AgNP (Ha et al., 2014; Nakashima et al., 2019). Dysregulation of autophagy, induced by pathway inhibition or excessive stimulation under external stressors, could explain these observations.

Moreover, Live cell Painting provided insights into the mechanisms of action (MOA) of drugs based on cell phenotypes. Amiodarone induced phospholipidosis in HepaRG cells by short-term exposure to the drug (Anthérieu et al., 2010), and alterations shown by AO staining in vesicle morphology and spatial distribution patterns (normalized moments, eccentricity) suggest that amiodarone might impact vesicular trafficking and dynamics (Supplemental Figure S10). Similarly, AO staining reveals significant insights into etoposide’s MOA by showing increased texture correlation and granularity in nuclei, indicative of the drug’s effect on nuclear morphology and its role in inducing DNA damage responses. These changes are consistent with etoposide’s mechanism as a topoisomerase II inhibitor, leading to cell cycle arrest and apoptosis (Montecucco et al., 2015). Conversely, despite its classification as safe for the liver (Class 0 in DILI severity), Orphenadrine’s impact on cellular structures and dynamics in Huh-7 cells reveals subtle effects through AO staining. This includes alterations in vesicle morphology and reduced measures of nucleoli and cytoplasmic mass displacement, suggesting potential changes in cellular protein synthesis and metabolic activity (Kornhuber et al., 1995; Rumore and Schlichting, 1985). Overall, Live Cell Painting enabled comprehensive assessments of cell health and morphological profiles under various treatment conditions, promising scalability for large-scale and cost-effective applications in the future.

Our results establish a live cell proof-of-concept framework for detecting intracellular events and profiling morphological changes using AO and high-content microscopy. While our framework successfully demonstrates its utility in assessing acidic vesicle dynamics and cellular morphological profiles, some limitations require consideration. For instance, we observed photobleaching in the AO-Green and AO-Red channels during repeated imaging sessions (10 consecutive acquisitions spaced 10 seconds apart), which suggests the need for improvements in AO’s photostability for extended kinetic studies (data not shown). Furthermore, caution is advised in interpreting AO staining, necessitating comparison with ground truth annotated compounds to validate new drug profiling. AO primarily assesses nucleic acids and acidic vesicles, thereby indirectly indicating alterations in numerous cellular signaling pathways. Moreover, optimizing AO concentrations to balance staining intensity and cytotoxicity remains a challenge across different cell lines, often requiring the integration of alternative stains like Hoechst to enhance contrast without compromising cell viability. Despite these challenges, our approach holds promise for broad applications across diverse cell types, diseases, and biological inquiries. Future studies should focus on evaluating AO’s efficacy in discerning varied phenotypes and mechanisms of action across a wider range of compounds, potentially integrating clinical samples and multi-omics data to enhance its clinical relevance and applicability.

## Conclusion

Live Cell Painting represents a straightforward assay for investigating live cells, offering an image-based profile and detailed insights into intracellular events. The Live Cell Painting captures acidic vesicle dynamics and cellular response to low concentrations of nanoparticles and generates hypotheses about the interaction between nanoparticles and cells. In addition, it is used to define groups of cells based on their phenotype after treatment with drugs that induce liver injury. Acridine orange is a suitable dye for image-based profiling, providing interesting information about the condition and health of cells while providing insights for new biological questions/research. We believe the assay could benefit the field of image-based live cell profiling and potentially be used in toxicology and drug development in the future.

## Material and Methods

### a. Cell culture

Huh-7 human hepatoma cells (No. JCRB0403), hereinafter referred to as Huh-7, were obtained from the Japanese Collection of Research Bioresources Cell Bank (JCRB, Japan). Cells were maintained in culture in Dulbecco’s Modified Eagle’s Medium (DMEM, Life Technologies, Canada) supplemented with 10% Fetal Bovine Serum (FBS, Gibco, South America) and 1% antibiotics (Penicillin/Streptomycin (Pen/Strep)) (Gibco, Grand Island - USA). Cells were incubated in a Panasonic incubator at 37°C, 95% humidity, and 5% CO_2_ and were passed as required for maintenance when confluency reached approximately 80%. Cultures were determined to be free of mycoplasma at each thaw using the direct DNA staining method (with Hoechst 33342 (Invitrogen, USA)) and inspected under fluorescence microscopy. All experiments were performed with cells free of mycoplasma.

### b. Acridine Orange staining

Huh-7 cells were plated at a density of 8000 cells per well in 96-well plates and grown in DMEM medium supplemented with 10% FBS and 1% Pen/Strep. After 24 hours, the cells were treated in triplicate with 100 µL of a range of Acridine Orange (AO) concentrations (2.5, 5, 10, 20, and 40 µM) diluted in a non-supplemented medium (serum-free). Then, cells were incubated for 10 minutes at 37°C and 5% CO_2_. After 10 minutes, the AO solutions were replaced by 100 µL of FluoroBrite DMEM (Gibco, USA). Fluorescence images were acquired on Cytation 5 Hybrid Multidetection Reader (BioTek Instruments, Inc., Winooski, VT, USA), from herein called Cytation 5, using the filter cubes from Biotek named GFP (excitation/emission = 469/525 nm) and Propidium Iodide (excitation/emission = 531/647 nm).

### c. ATP viability test

We chose to assess cell viability by quantifying ATP (adenosine triphosphate) using the luminescence signal because AO interferes with common viability assays such as MTT, neutral red, and calcein AM. The ATP viability assay was performed according to the manufacturer’s instructions. Briefly, CellTiter-Glo® Buffer and lyophilized CellTiter-Glo® Substrate (Promega, USA) were placed in the bead bath to reach room temperature. Subsequently, the entire content of CellTiter-Glo® Buffer was mixed with the lyophilized CellTiter-Glo® Substrate. The mixture was vortexed to obtain a homogeneous solution, forming the CellTiter-Glo® Reagent. Huh-7 cells were plated at a density of 8000 cells per well in 96-well plates and grown in DMEM medium supplemented with 10% FBS and 1% Pen/Strep. The cells were treated in triplicate with 100 µL of a range of AO concentrations (2.5, 5, 10, 20, and 40 µM) diluted in non-supplemented medium (serum-free) and 100 µL of non-supplemented medium only (negative control), and incubated for 4 hours at 37°C and 5% CO_2_. After 4 hours of treatment with AO, 100 µL of CellTiter-Glo® Reagent was added on top of the treatments. The plate was shaken for 2 minutes on an orbital shaker to induce cell lysis. Subsequently, the contents of the plate were transferred in the same order to an opaque white 96-well plate (suitable for luminescence reading). Then, the plate was incubated at room temperature for 10 minutes to stabilize the luminescent signal. Subsequently, the luminescence of the wells was read in Cytation 5. The luminescence value was normalized as a percentage to the luminescence value in wells treated only with culture medium (which was considered 100%).

### d. Acidification inhibitors assay

Huh-7 cells were plated into a 96-well plate in an 8000 cells/well density. After 24 hours of incubation in a Panasonic incubator at 37°C, 95% humidity, and 5% CO_2_, the cells were treated with vesicle acidification inhibitors Chloroquine 50 μM (Cayman Chemical Company) and Bafilomycin A 1.5 μM (Cayman Chemical Company) mixed with either Acridine Orange 20 μM or Lysotracker Deep Red (ThermoFisher) at 75 nM. Cells were treated for 0.5 hours and 4 hours, then the treatment was removed and FluoroBrite DMEM (Gibco, USA) was added for image acquisition. Sixteen sites were acquired per well using 20x Olympus objective plan fluorite NA 0.45 (1320517) with Cytation 5 Hybrid Multidetection Reader (BioTek Instruments, Inc., Winooski, VT, USA). Images from Lysotracker Deep Red were acquired using CY5 led cube (excitation/emission = 628/685 nm), and from AO-treated cells using GFP (excitation/emission = 469/525 nm) and Propidium Iodide (PI) (excitation/emission = 531/647 nm) filters. The total intensity was calculated by multiplying the “Cells_Intensity_IntegratedIntensity_AO-Red”, a feature that represents the total pixel intensity in each cell in the AO-Red, by the total number of cells analyzed. The radial distribution presented was calculated by dividing the cells into 4 concentric rings and measuring the intensity of PI in each concentric ring.

### e. Dose-response assay for silver nanoparticles

Huh-7 cells were plated in an 8000 cells/well density in a 96-well plate and incubated in a Panasonic incubator at 37 °C, 95% humidity, and 5% CO_2_. After 24 hours, cells were treated with silver nanoparticles (AgNP) of size 40 nm (Ag40) or 100 nm (Ag100) (Sigma-Aldrich, 730807-25ML and 730777-25ML). The concentrations tested for Ag100 were 0.009, 0.017, 0.035, 0.069, 0.139, 0.278, 0.556, and 1.111 μM, and for Ag40 were 0.0011, 0.0022, 0.0043, 0.0087, 0.0174, 0.0347, 0.0694, 0.1389 μM. Cells were treated for 24 hours with the concentrations stated. Then, AO 20 μM was added for 10 minutes, and the media was replaced by 100 μL of FluoroBrite DMEM (Gibco, USA) for image acquisition. The same test was done in parallel to assess the cell viability using MTT at 5 ug/mL (3-(4,5-Dimethylthiazol-2-yl)-2,5-Diphenyltetrazolium Bromide) (Invitrogen). Sixteen sites per well were acquired using a 20x Olympus objective plan fluorite NA 0.45 (1320517) in Cytation 5 Hybrid Multidetection Reader (BioTek Instruments, Inc., Winooski, VT, USA), with the incubator on to maintain 37 °C and 5% CO_2_. Images from AO were acquired using GFP (excitation/emission = 469/525 nm) and Propidium Iodide (PI) (excitation/emission = 531/647 nm) filters. Sixteen sites per well were acquired, and data was processed and analyzed according to image processing and profiling described below.

### f. Drug screening assay

Huh-7 cells were plated at a cell density of 8000 cells per well in a 96-well plate. The cell suspension was carefully added to each well, ensuring uniform distribution. The plated cells were allowed to sediment at the bottom of the wells by leaving the plate undisturbed inside the hood for 20 minutes. Subsequently, the plate was transferred to the cell culture incubator and left undisturbed until the following day to allow cell adhesion and growth. 24 hours later, the compounds were prepared by diluting them to concentrations of 1 μM and 10 μM using an incomplete DMEM medium. The cells were treated with the respective compound solutions (Table 1) for 24 hours. Each replicate was performed on different days, and unique plate layouts were designed for each day to account for potential day-to-day variations. On the next day, the culture medium was aspirated, and subsequently, 100 μL of acridine orange (AO) solution at 20 μM concentration, was added to each well for cell staining. After a 10-minute incubation, the AO solution was aspirated, and 100 μL of FluoroBrite DMEM (Gibco, USA) medium was added to each well. Then, 25 sites per well were acquired using a 20x Olympus objective plan fluorite NA 0.45 (1320517) in the fluorescence microscope Cytation 5 Hybrid Multidetection Reader (BioTek Instruments, Inc., Winooski, VT, USA), on AO-Green and AO-Red channels.

### g. Image processing and analysis

A cellpose model was trained on images from Huh-7 cells stained with AO to identify the nuclei object using the cellpose GUI (Pachitariu and Stringer, 2022) (model available here https://github.com/broadinstitute/scripts_notebooks_fossa/tree/main/cellpose/Acridine_Orange_Model_Huh7). One image from every experiment was randomly selected to create the train and test. The models were used to segment the nuclei on all our analysis pipelines using the CellProfiler plugin RunCellpose which runs cellpose custom models (Weisbart et al., 2023). Then a pipeline was created on CellProfiler version 4.2.5 to segment the nuclei, cells, cytoplasm, nucleoli, and vesicles, and an overlay of the segmented objects was created above the images to check for image segmentation (Carpenter et al., 2006; Stirling et al., 2021). The pipeline was run for one image of every well in every plate before running the analysis, to check for errors in object segmentation. This step of checking object segmentation joined with the cellpose model used, was considered the quality control for all assays performed. An example pipeline is available here https://github.com/broadinstitute/scripts_notebooks_fossa/tree/main/cellprofiler/LiveCellPainting_pipeline.

After segmentation, features were extracted from the single-cell objects using CellProfiler modules. Features related to the area and shape of the objects were extracted, such as area, perimeter, and HuMoment (Hu, 1962); the integrated, mean, and other features related to pixel intensity were measured in every channel and object (cell, nuclei, and cytoplasm); the degree and nature of textures in images and objects to quantify their roughness and smoothness (Haralick et al., 1973). The cells were segmented into 4 concentric rings going from the center of the cell to the border of the cell, and the total and mean intensity of the AO-Green and AO-Red was measured within each ring. The pipelines were run using a version of CellProfiler optimized to run inside Amazon Web Services, the Distributed-CellProfiler (Weisbart and Cimini, 2023). To join the CSV files generated per site, the pycytominer function collate from the “cyto utilities” package was used (Serrano et al., 2023).

### h. Profiling

To process the features and obtain a meaningful profile (Chandrasekaran et al., 2021), Pycytominer was used to process image-based profiles (Serrano et al., 2023). Profiles were aggregated from single-cell to well-aggregated profiles using the median function from the Pycytominer function “aggregate”. Annotation was also performed with Pycytominer to add all relevant information for each dataframe such as compound name, the concentration used, cell type, etc. The features were normalized relative to the negative control on a per-plate basis to address batch effects using the RobustMAD function, which takes the feature value (x) minus the median divided by the median absolute value (scaled feature = (x - median/mad). Feature selection was performed to exclude features that have more than 5% of NA values, features with low variance, features that correlate higher than 90%, and features that have values higher than 500 (outliers). Notebooks for profiling are available here https://github.com/broadinstitute/scripts_notebooks_fossa/tree/main/profiles/cellprofiler. Batch correction was applied to correct the effect of performing the experiment on different days. pyCombat (an implementation of ComBat in Python; example notebook is available here https://github.com/broadinstitute/scripts_notebooks_fossa/blob/main/pycombat_umap/apply_pycombat_visualize.ipynb) (Behdenna et al., 2020) was used because of its characteristic of correcting for batch effect in small sample sizes. From herein, when a profile is defined, we consider that all these steps were applied before analyzing the results.

### i. Profile Evaluation

The replicate retrieval ability was measured using mean Average Precision. It measures the ability of a replicate to retrieve other replicates without classifying the negative control as a replicate. A query is created to retrieve replicates or negative control, and a rank order is created. From those, the area under the precision-recall curve is measured. More details are available in (Dijk et al., 2023). The test was performed using the evalzoo/matric library available at https://github.com/cytomining/evalzoo/tree/main/matric.

To visualize the profiles in a dimensionality-reduced space, t-SNE (t-distributed stochastic neighbor embedding) (Maaten and Hinton, 2008) or UMAP (Uniform Manifold Approximation and Projection) (McInnes et al., 2020) were used to generate the vectors for visualization. The embeddings were generated for 100 iterations, and the mean and standard deviation embedding were calculated and presented in the plots.

HDBSCAN (Hierarchical Density-Based Spatial Clustering of Applications with Noise) was performed by following the example available at https://umap-learn.readthedocs.io/en/latest/clustering.html. UMAP is used for dimensionality reduction, and HDBSCAN is performed to predict the labels based only on the density of the points (unsupervised clustering method).

### j. Statistical analysis

Statistical analysis was performed to compare the negative control to each compound presented on every assay. The statannotations library was used for all the tests performed in this paper (Charlier et al., 2022). In the acidification inhibitors assay (Figure 2) independent t-test was used to test the total intensity differences between the negative control and acidification inhibitors. In the dose-response assay with AgNP (Figure 3), Mann-Whitney was used to test all concentrations of AgNP against the negative control. More details for each test can be found in the caption of each plot, but in general, the p-value was displayed as *: 0.01 < p <= 0.05, **: 0.001 < p <= 0.01, ***: 0.0001 < p <= 0.001, and ****: p <= 0.0001. An example notebook is available here https://github.com/broadinstitute/scripts_notebooks_fossa/blob/main/individual_feature_and_statistics/Individual_Features_Statistics.ipynb.

## Data Availability Statement

Data available on request from the authors.

## Acknowledgments

Funding was provided by São Paulo Research Foundation (FAPESP) #2022/01483-4, #2019/24033-1, #2023/06143-0 and #2020/01218-3.

Funding was also provided by the National Institutes of Health (NIH COBA P41 GM135019 to BAC and AEC and MIRA R35 GM122547 to AEC). This project has been made possible in part by grant number 2020-225720 to BAC from the Chan Zuckerberg Initiative DAF, an advised fund of the Silicon Valley Community Foundation.

## Supplementary material

**Supplemental Figure 1.**
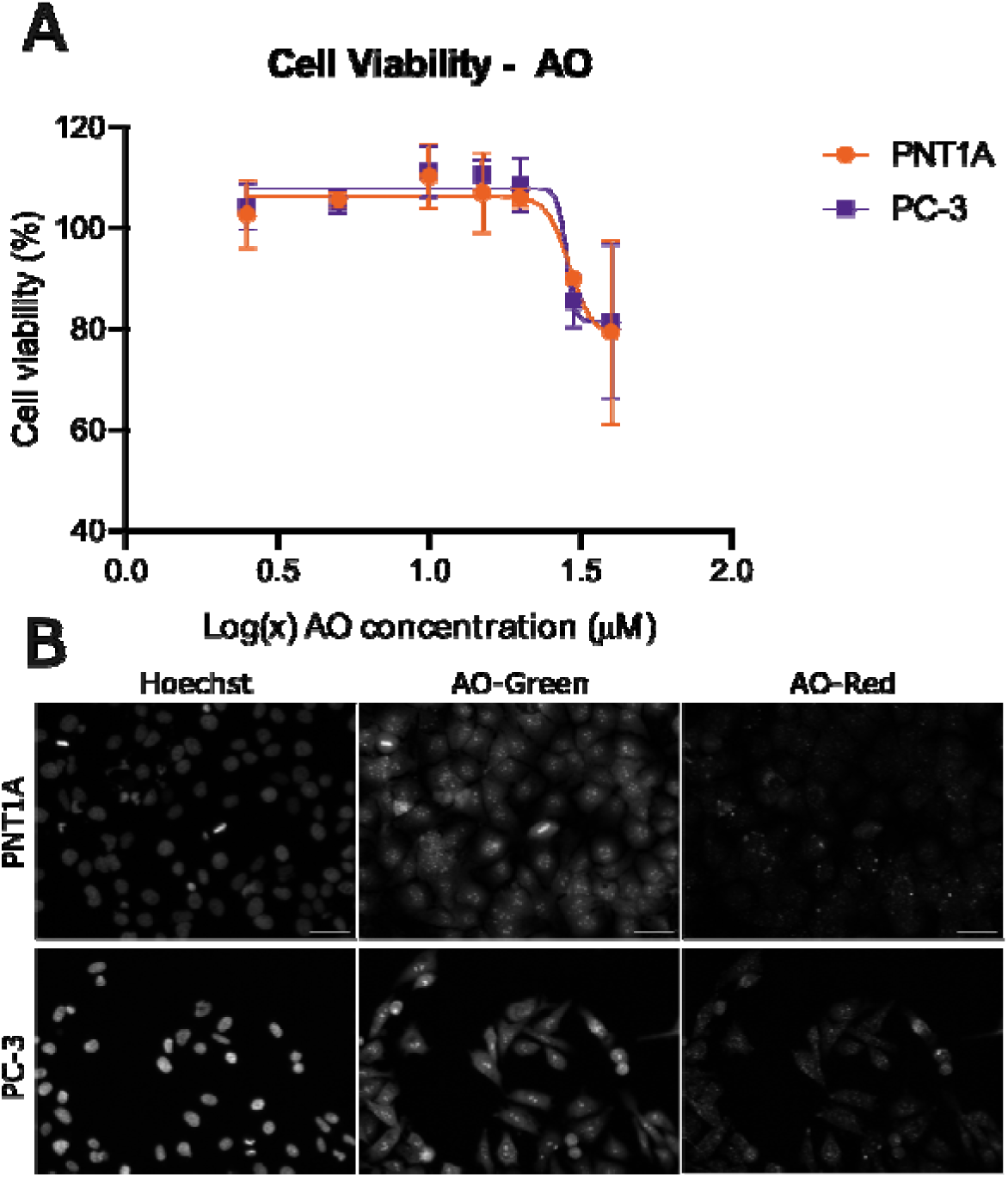
Cell viability of PNT1A and PC-3 cells in the presence of AO assessed by ATP concentration. Cells were treated with AO (2.5, 5.0, 10.0, 20.0, and 40.0 μM) for 4 hours. Then, viability was assessed using CellTiter-Glo® by measuring ATP concentration with luminescence and images were acquired by Cytation 5, Biotek (20x objective). ATP concentration on the cells was determined by fitting a linear function to the ATP standard curve and calculating the relative viability compared to the negative control. (A) Viability curve for PNT1A and PC3 cells exposed to AO. (B) Representative images for PNT1A and PC-3 cells exposed to 2.5 μM of AO, respectively, represented by the Hoechst, AO-Green and AO-Red channels. The lowest concentration tested, 2.5 μM, was selected as it exhibited no cytotoxicity in either cell line. Higher concentrations than 2.5 μM do not decrease cell viability as seen in (A), but cell morphology was visually different than the control. Due to the low concentration of AO, differentiation between the nucleus and cytoplasm was less distinct, cells were stained with Hoechst to facilitate the nuclei segmentation during image analysis. Nonetheless, structures such as the nucleus, nucleoli, vesicles, and cytoplasm were still visible in both cell lines. Optimization of AO concentration should be performed for each cell line. Scale bar = 50 μm. Statistical test performed with two-way ANOVA followed by Sidak’s multiple comparisons test: * p < 0.05.

**Supplemental Figure 2.**
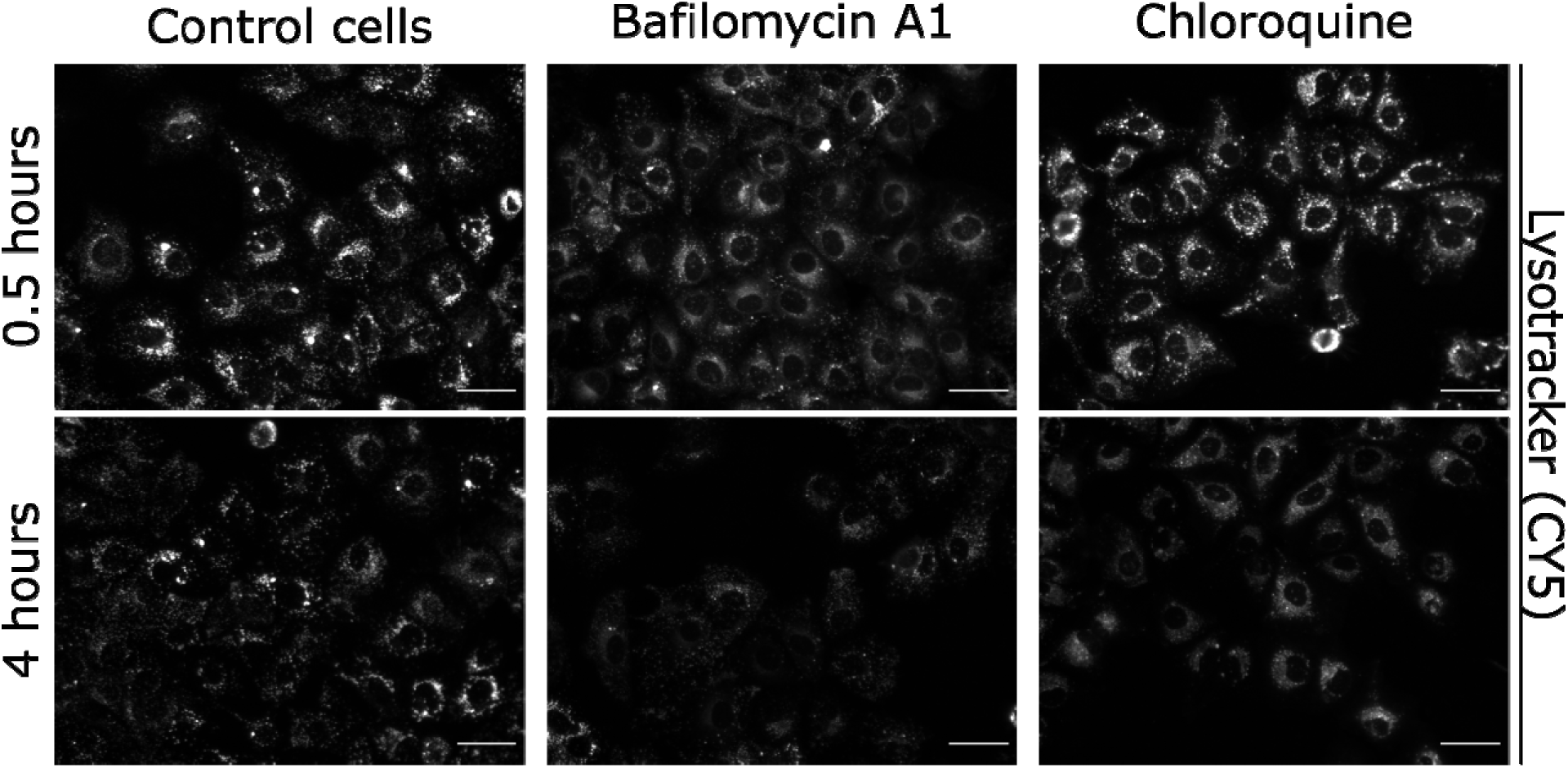
Validation of acidification inhibitors using Lysotracker Deep Red. Huh-7 cells were plated in a 96-well plate at a density of 8000 cells/well. The following day, the cell were treated with solutions containing the inhibitors Bafilomycin A1 1.5 µM or Chloroquine 50 µM added by Lysotracker 75 nM for 0.5 h and 4 h. The images were acquired on the Cytation 5 Hybrid Multidetection Reader (BioTek Instruments, Inc., Winooski, VT, USA using 20× objective. Representative images of the treatments with Lysotracker + inhibitors, acquired in the CY5 channel, represented in grayscale. Scale = 50 µm.

**Supplemental Figure 3.**
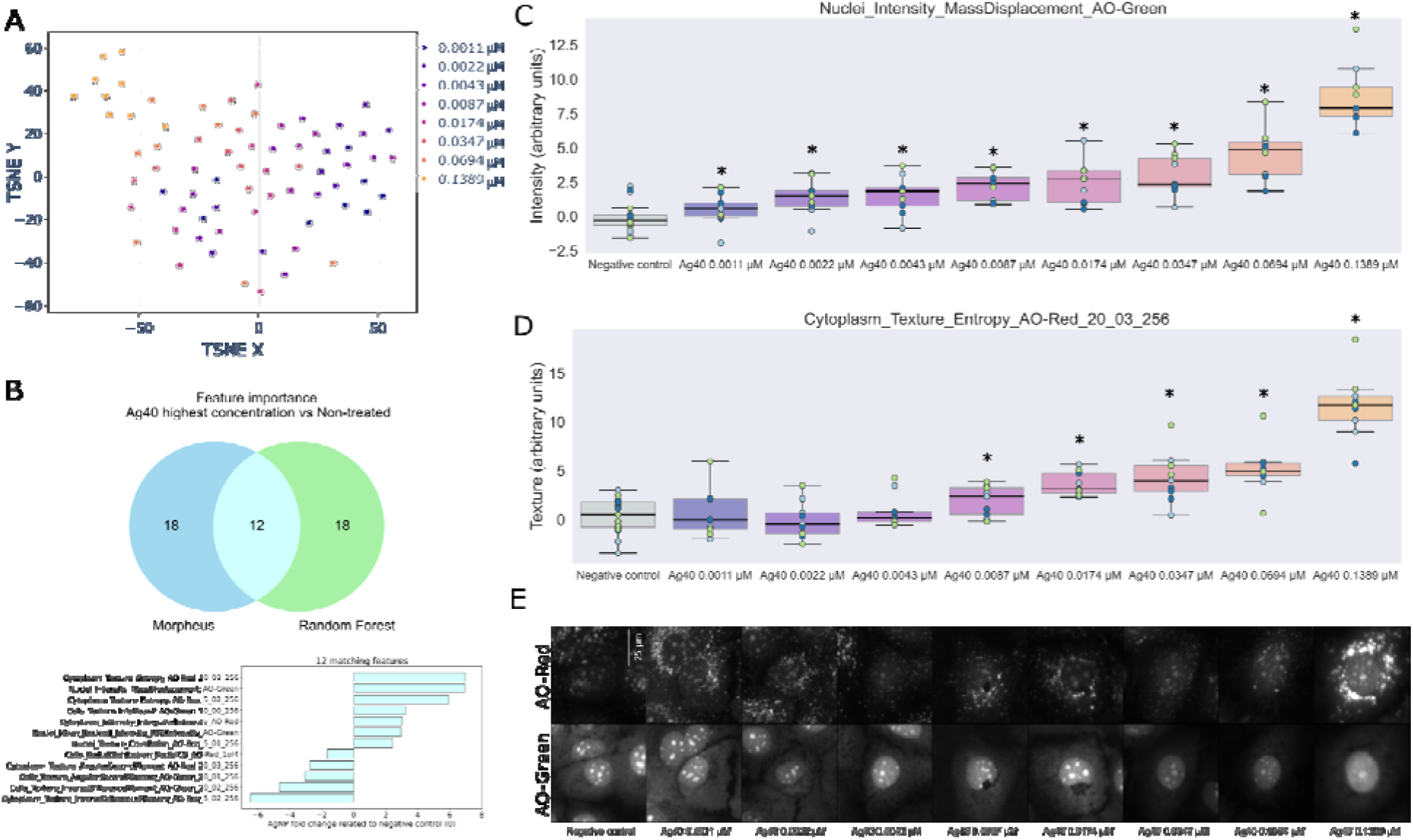
The Live Cell Painting assay detects the dose-response effect on Huh-7 cells induced by silver nanoparticles (AgNP). Huh-7 cells were treated with Ag100 (nanoparticle size of 40 nm) for 24 hours with increasing concentrations of nanoparticles. Then the image were acquired after AO staining (Cytation 5, 20x objective). Objects were segmented using cellpose and features were extracted using CellProfiler; next, features were aggregated by well, normalized, and feature selected using pycytominer, and batch corrected with pyComBat. (A) t-SNE was used for dimensionality reduction and visualization. After 100 iterations, the mean and standard deviation embedding were calculated; (B) Random forest ran for 100 iterations to differentiate between the classes: negative control and Ag40 0.0694 and 0.1389 μM. Then, the mean feature importance and standard deviation were calculated. Morpheus software was used to perform marker selection between the same groups classified in Random Forest. The 30 most important features were selected and compared. Among those, 8 features matched between the methods and were plotted to show the fold change relative to the negative control. The two features highlighted with black boxes are presented in C and D; (C) The intensity of AO-Green in the nuclei; (D) The spatial moment of the nuclei, representing shape, size, rotation, and location of the nuclei object, and (E) Representative images were retrieved from the dataset using k-means clustering (Garcia-Fossa et al., 2023). Experiments were performed in thre independent replicates, n =3 and each point in the plots is well-aggregated data, and points ar colored by the batch in C and D. Independent t-test calculated using the statannotations library. * p statistical difference < 0.05.

**Supplemental Figure 4.**
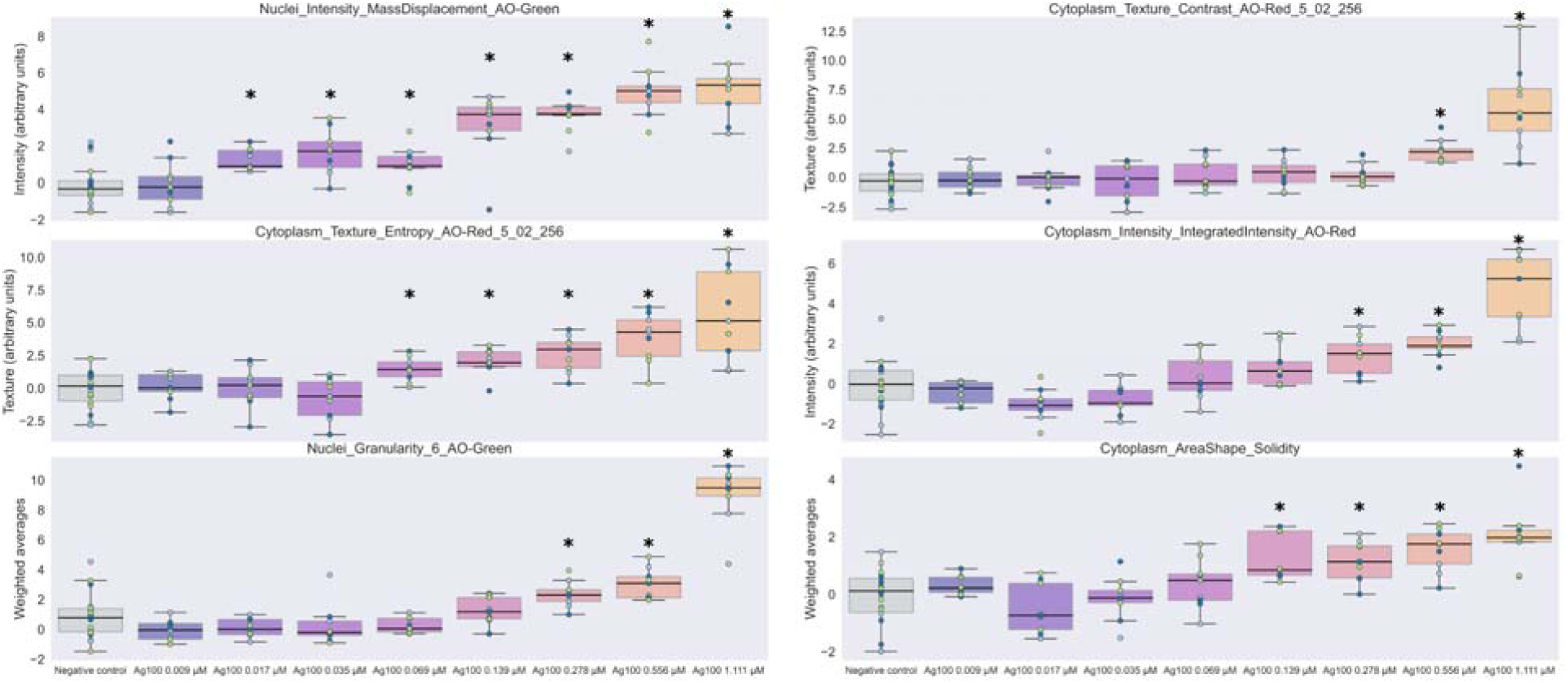
Features that have a dose-response behavior to Ag100 treatment, all showing an increase in overall values relative to the control. The six consistent features identified by both Morpheus and random forest analyses are plotted to illustrate fold changes relative to the negative control. These features include: nuclei intensity mass displacement on the AO-Green channel (movement of mass within nuclei); cytoplasm texture contrast on the AO-Red channel (variation in texture within acidic vesicles); cytoplasm texture entropy on the AO-Red channel (randomness in acidic vesicles texture patterns); cytoplasm intensity integrated intensity on the AO-Red channel (overall fluorescence intensity within acidic vesicles in cytoplasm); nuclei granularity on the AO-Green channel (measurement of uniformity and distribution of texture elements); and cytoplasm area shape solidity (compactness of cytoplasm shape).* p statistical difference < 0.05.

**Supplemental Figure 5.**
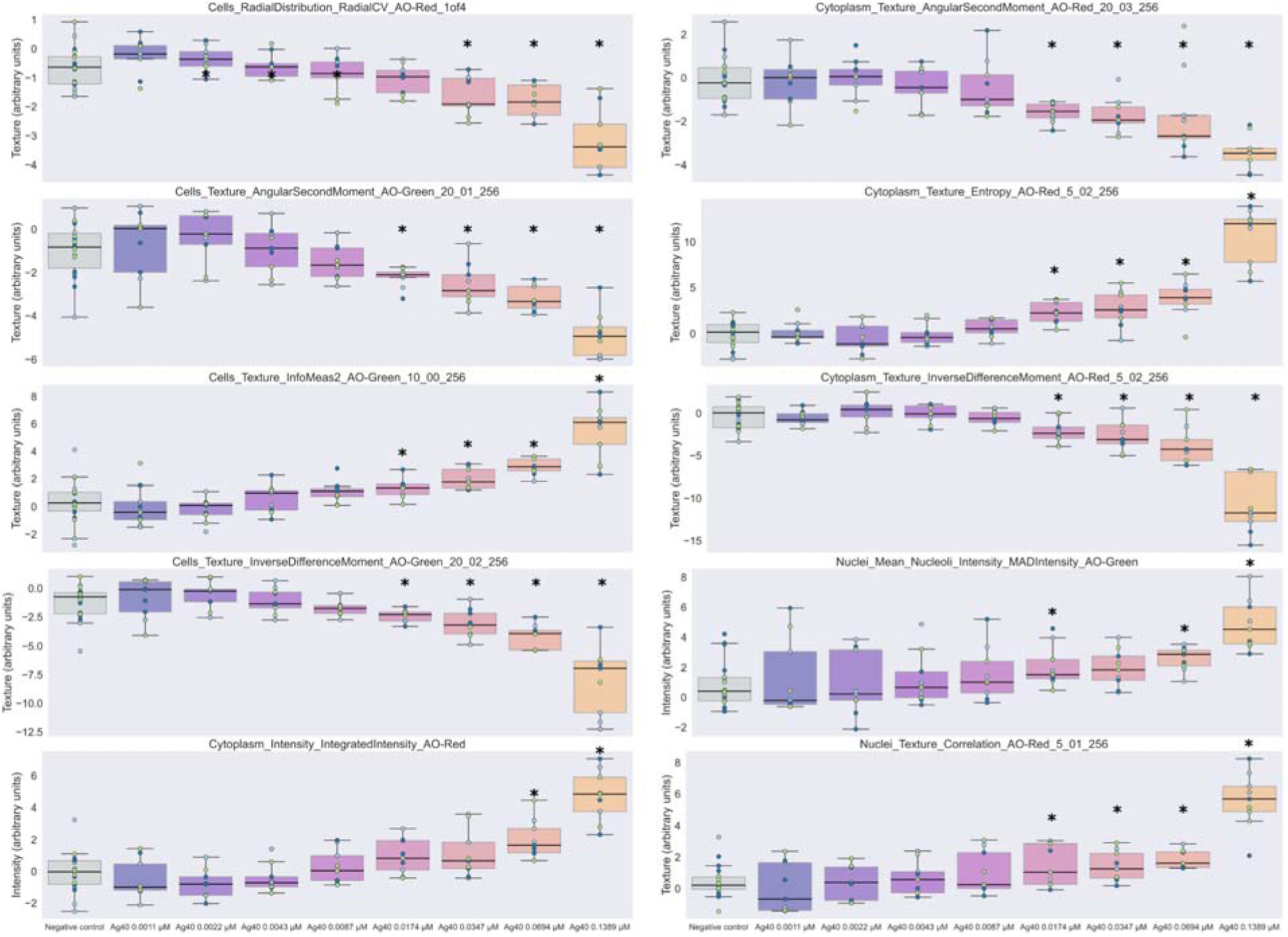
The ten features identified by random forest and Morpheus that have a dose-response behavior to Ag40 treatment, are plotted to illustrate fold changes relative to the negative control. These features include: cells radial distribution on the AO-Red channel (distribution pattern of acidic vesicles within the cell); cytoplasm texture angular second moment on the AO-Red channel (local homogeneity of acid vesicles in cytoplasm texture in angular patterns); cell texture angular second moment on the AO-Green channel (local homogeneity of cell texture in angular patterns); cytoplasm texture entropy on the AO-Red channel (randomness in cytoplasmatic acidic vesicles texture patterns); cells texture infomeas2 on the AO-Green channel (measure of texture complexity in cells); cytoplasm texture inverse difference moment on the AO-Red channel (local homogeneity of cytoplasmatic acidic vesicles texture in pixel intensity differences); cells texture inverse difference moment on the AO-Red channel (local homogeneity of acidic vesicles texture in pixel intensity differences); nuclei mean nucleoli intensity mad intensity on the AO-Green channel (average intensity of nucleoli within nuclei); cytoplasm intensity integrated intensity on the AO-Red channel (overall acidic vesicles fluorescence intensity within the cytoplasm); nuclei texture correlation on the AO-Red channel (correlation between texture elements within nuclei).* p statistical difference < 0.05.

**Supplemental Figure 6.**
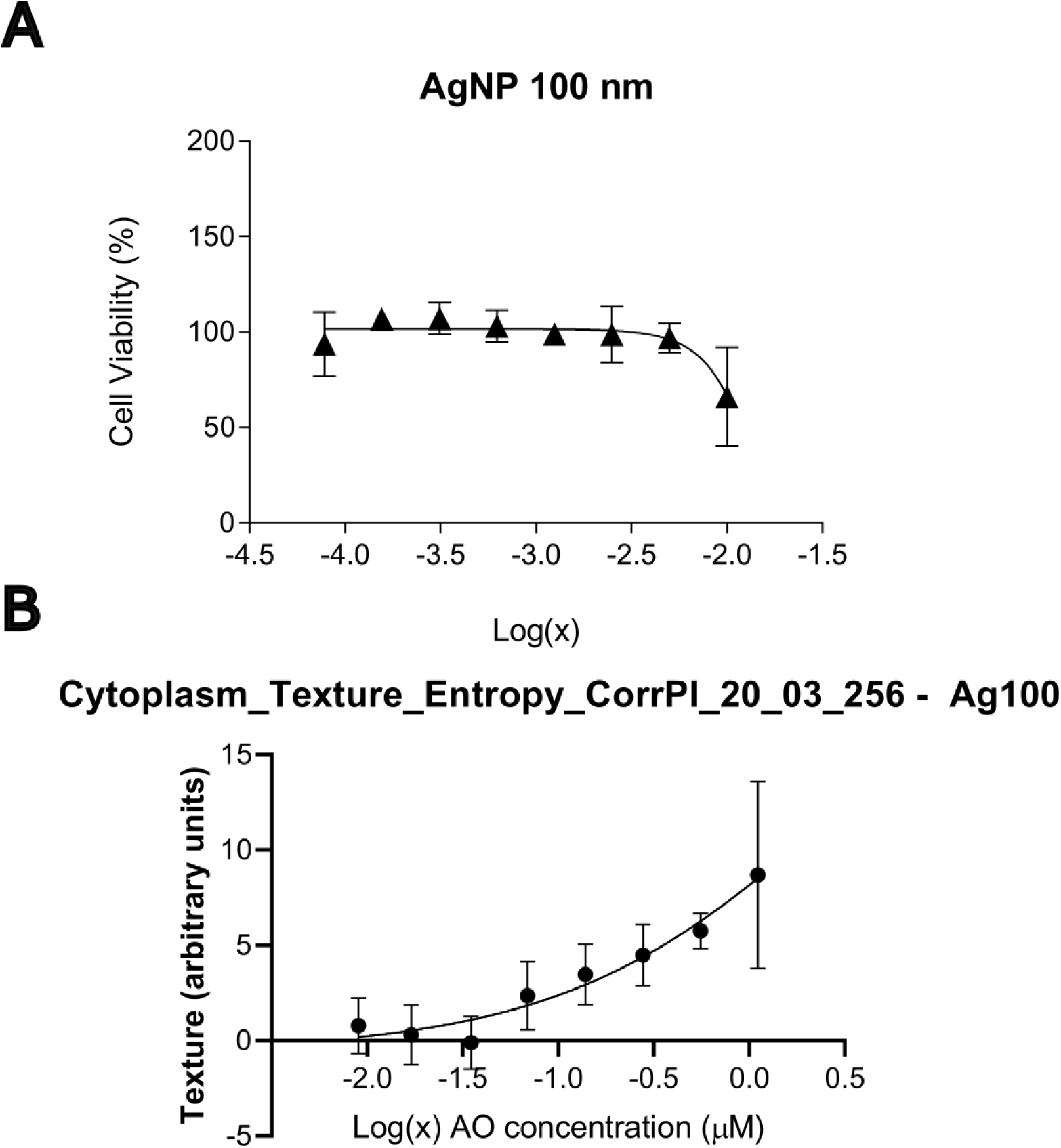
Comparison of IC50 calculation using MTT data or feature data. (A) The cells were treated with enhancing concentrations of AgNP 100 nm for 24 hours. Then, MTT 5 mg/mL was added and incubated for 2 hours. MTT absorbance was read on 570 nm using Cytation 5 Biotek. (B) The feature median values for each AgNP concentration were used to calculate the IC50.

**Supplemental Figure 7.**
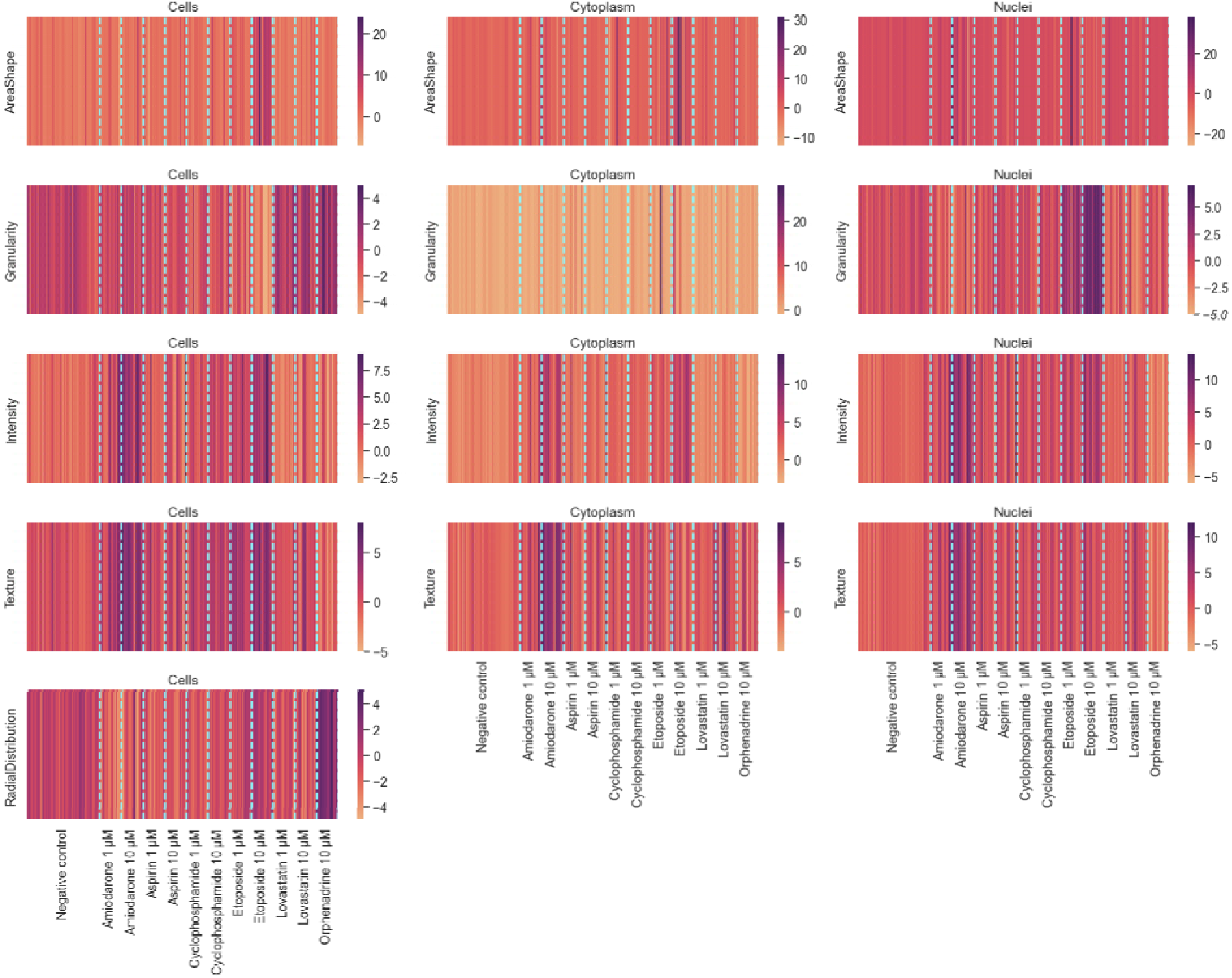
Features extracted from DILI-drugs assay after feature selection and split by categories and compartments. For the features in each category and each compartment, we calculated the principal component 1 to be a summary of all the features in that category. All features were normalized to the negative control, which has zero value in this matrix. The treatments induce a variation for each category that can be lower or higher than the negative control.

**Supplemental Figure 8.**
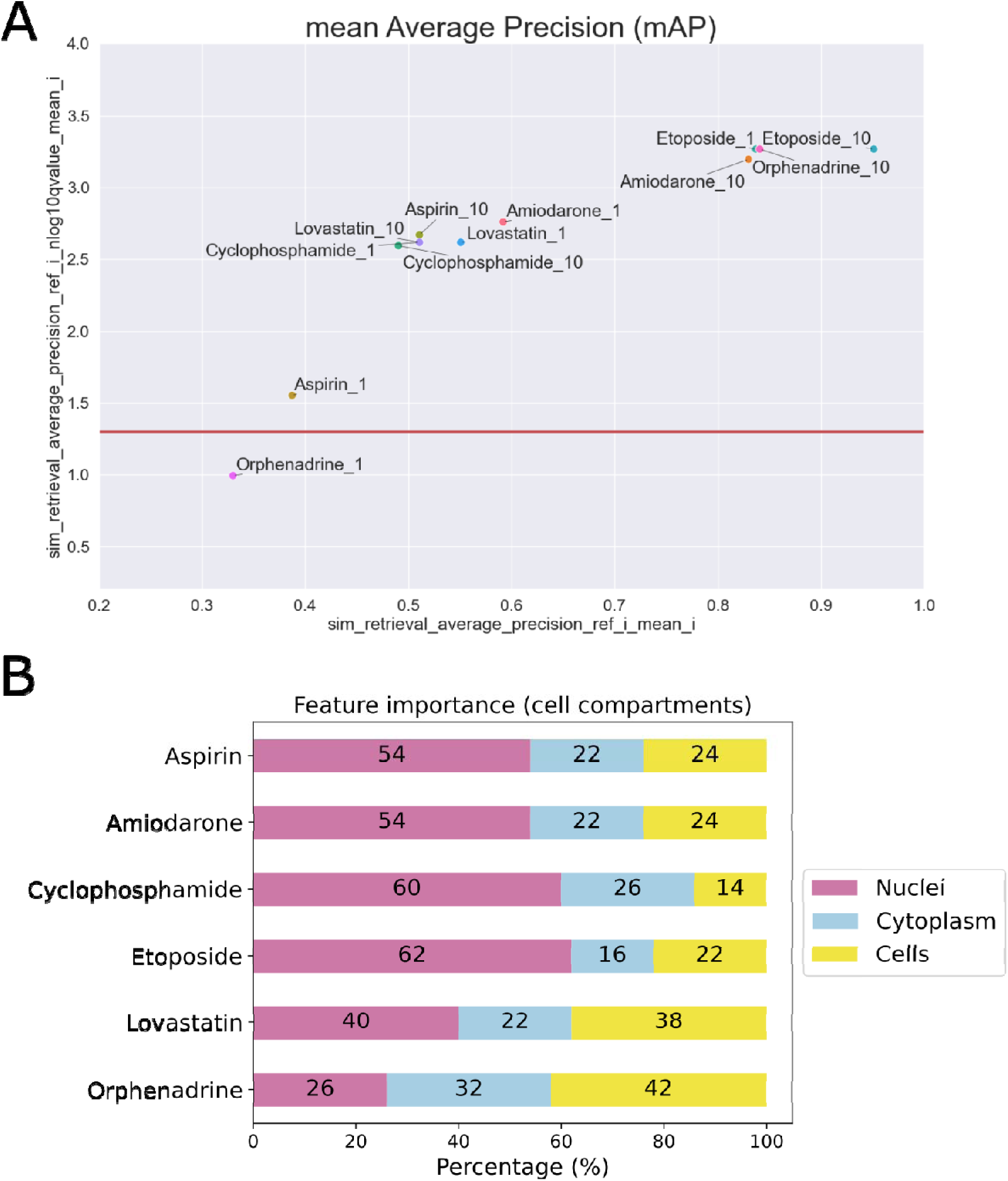
Evaluation of technical replicability of the profiles and compartments’ contribution for the 30 most important features. (A) Retrievable phenotypes are above the threshold (red line in x-axis, significance threshold 0.05). These compounds’ ability to retrieve themselves relative to the negative control is defined by their mean Average Precision (mAP) in the x-axis, and the q-value in the y-axis helps define the threshold of 0.05; (B) The contribution of each compartment for the most important features: Nuclei, Cytoplasm, and Cells.

**Supplemental Figure 9.**
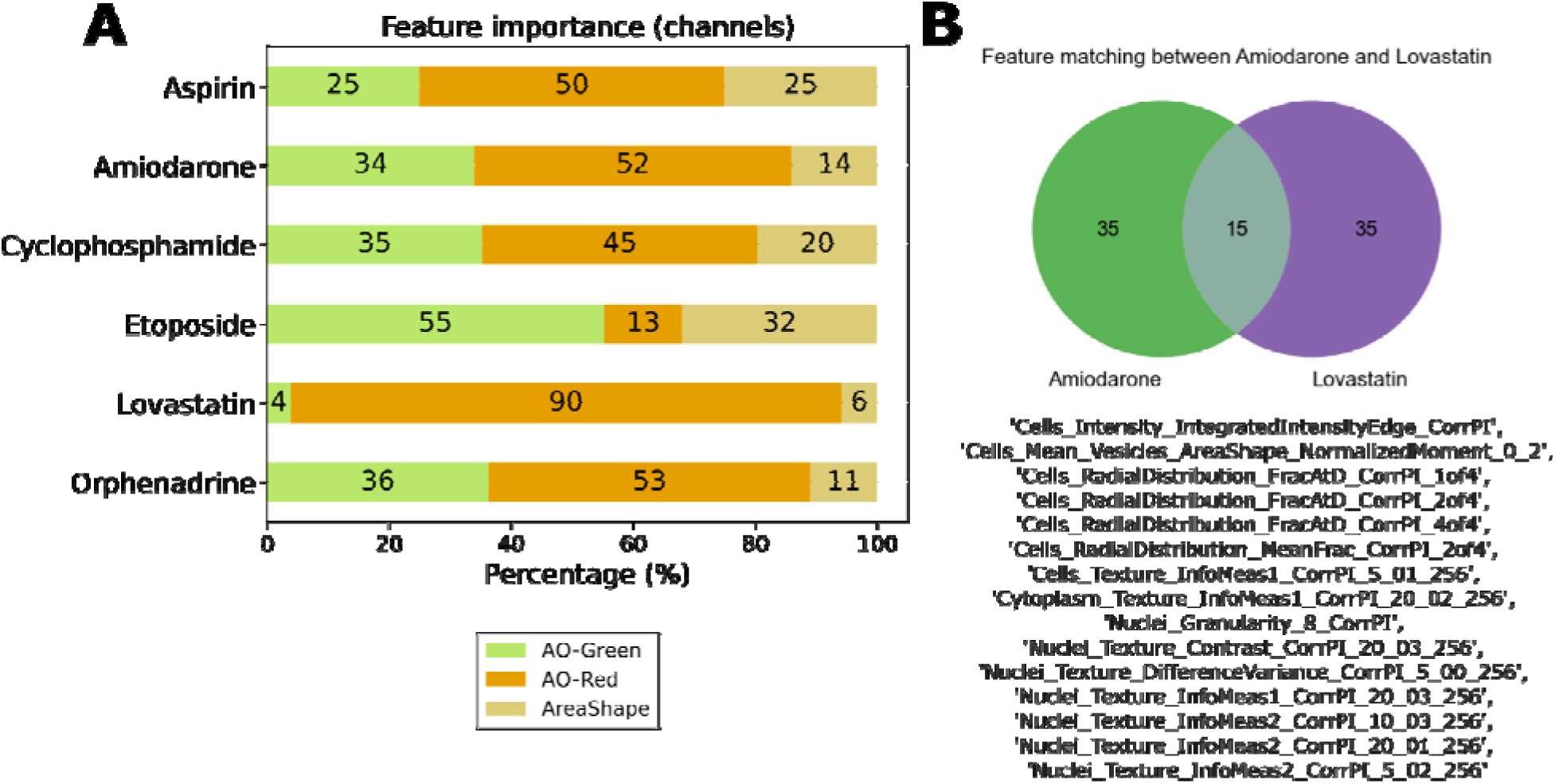
Feature matching between amiodarone and lovastatin. After selecting the 50 most significant features using Random Forest and Morpheus software, the matching features between the compounds was obtained. (A) The percentage of the 50 most important features that belong to AO-Green, AO-Red or AreaShape channels. For lovastatin and amiodarone, the percentage of features associated with the AO-Red are high. (B) All matching features are related to the AO-Red channel except for one associated with the shape of the vesicles.

**Supplemental Figure 10.**
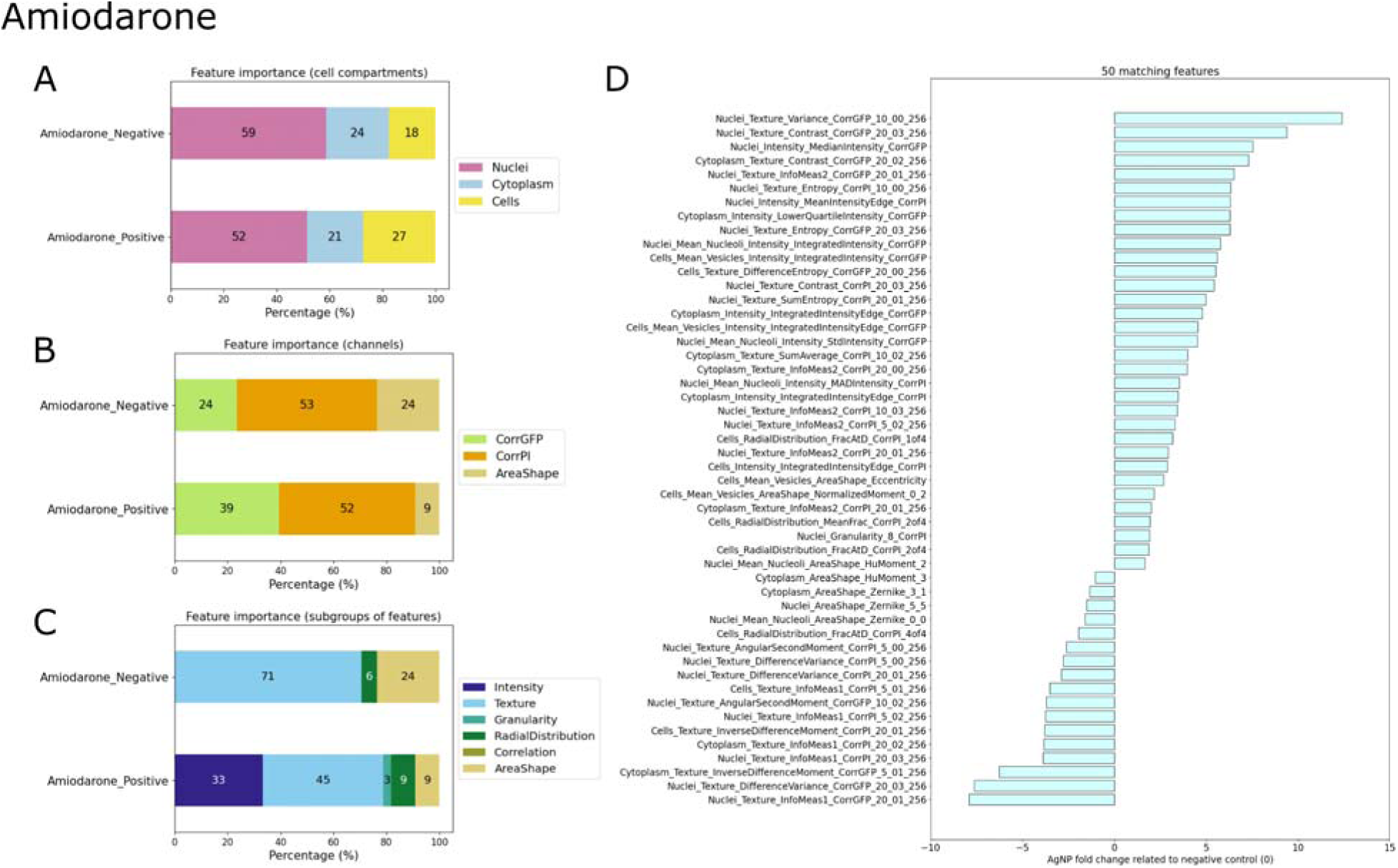
Key features defining amiodarone clustering. After selecting the 50 most significant features using Random Forest and Morpheus software, the median value of each feature was subtracted from the median value of the negative control. Features were then categorized as either negative or positive relative to the negative control. The results are presented as follows: (A) cell compartments, (B) channels, (C) subgroups of features, and (D) the complete list of features along with their fold changes relative to the negative control.

**Supplemental Figure 11.**
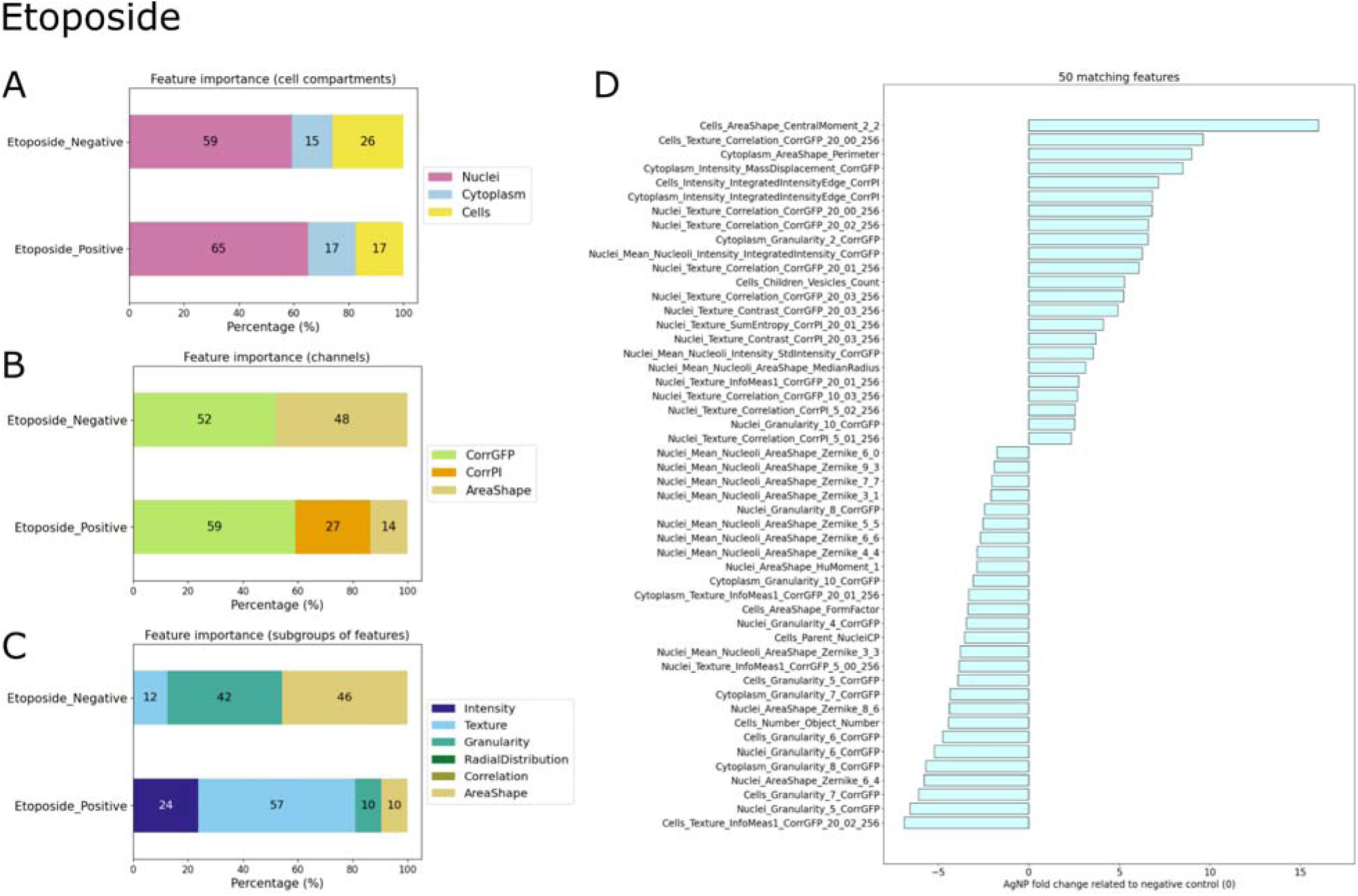
Key features defining etoposide clustering. After selecting the 50 most significant features using Random Forest and Morpheus software, the median value of each feature was subtracted from the median value of the negative control. Features were then categorized as either negative or positive relative to the negative control. The results are presented as follows: (A) cell compartments, (B) channels, (C) subgroups of features, and (D) the complete list of features along with their fold changes relative to the negative control.

**Supplemental Figure 12.**
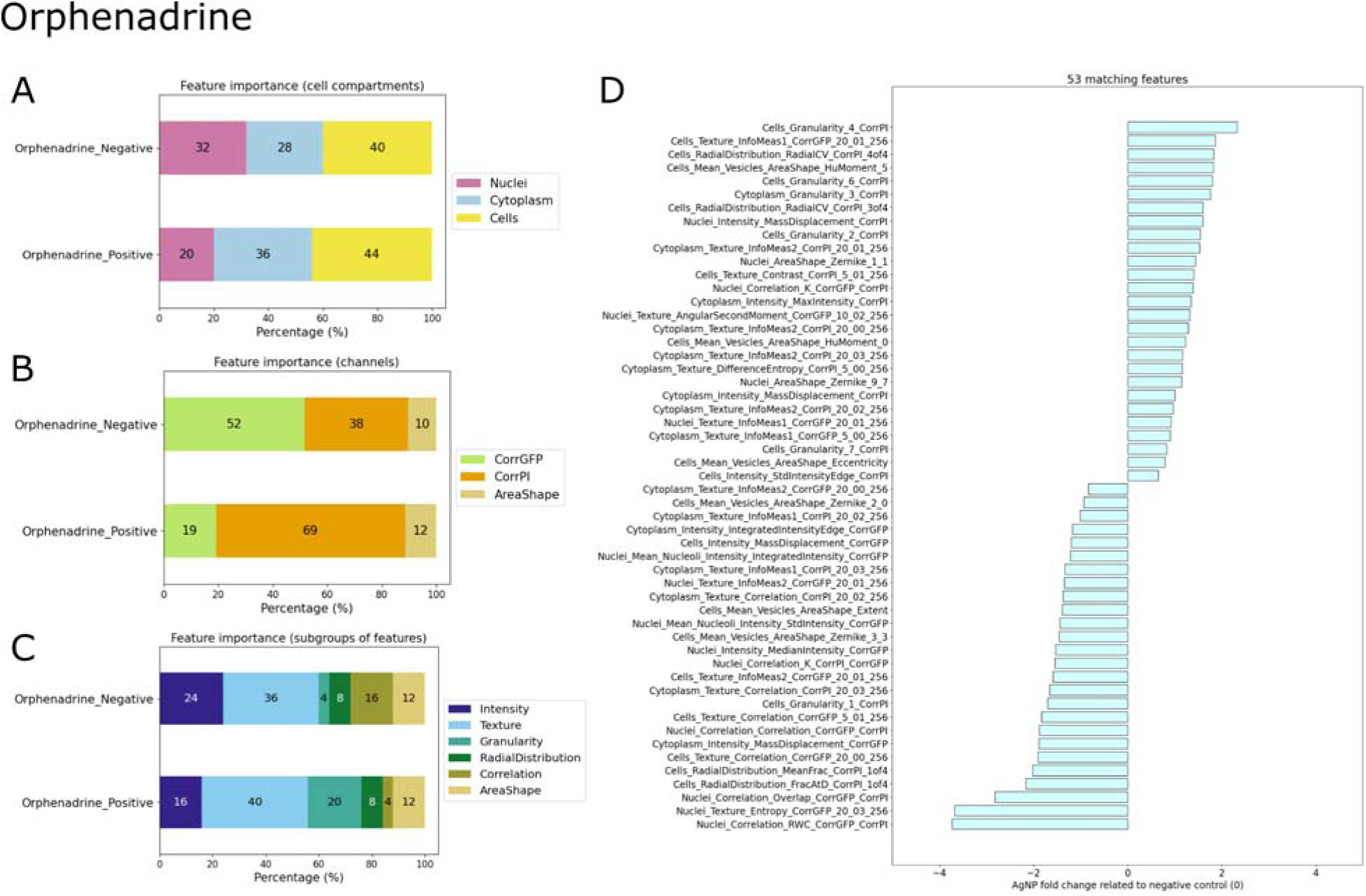
Key features defining orphenadrine clustering. After selecting the 50 most significant features using Random Forest and Morpheus software, the median value of each feature was subtracted from the median value of the negative control. Features were then categorized as either negative or positive relative to the negative control. The results are presented as follows: (A) cell compartments, (B) channels, (C) subgroups of features, and (D) the complete list of features along with their fold changes relative to the negative control.

