## Supplemental Material for "Live Cell Painting: image-based profiling in live cells using Acridine Orange"

### Supplementary material

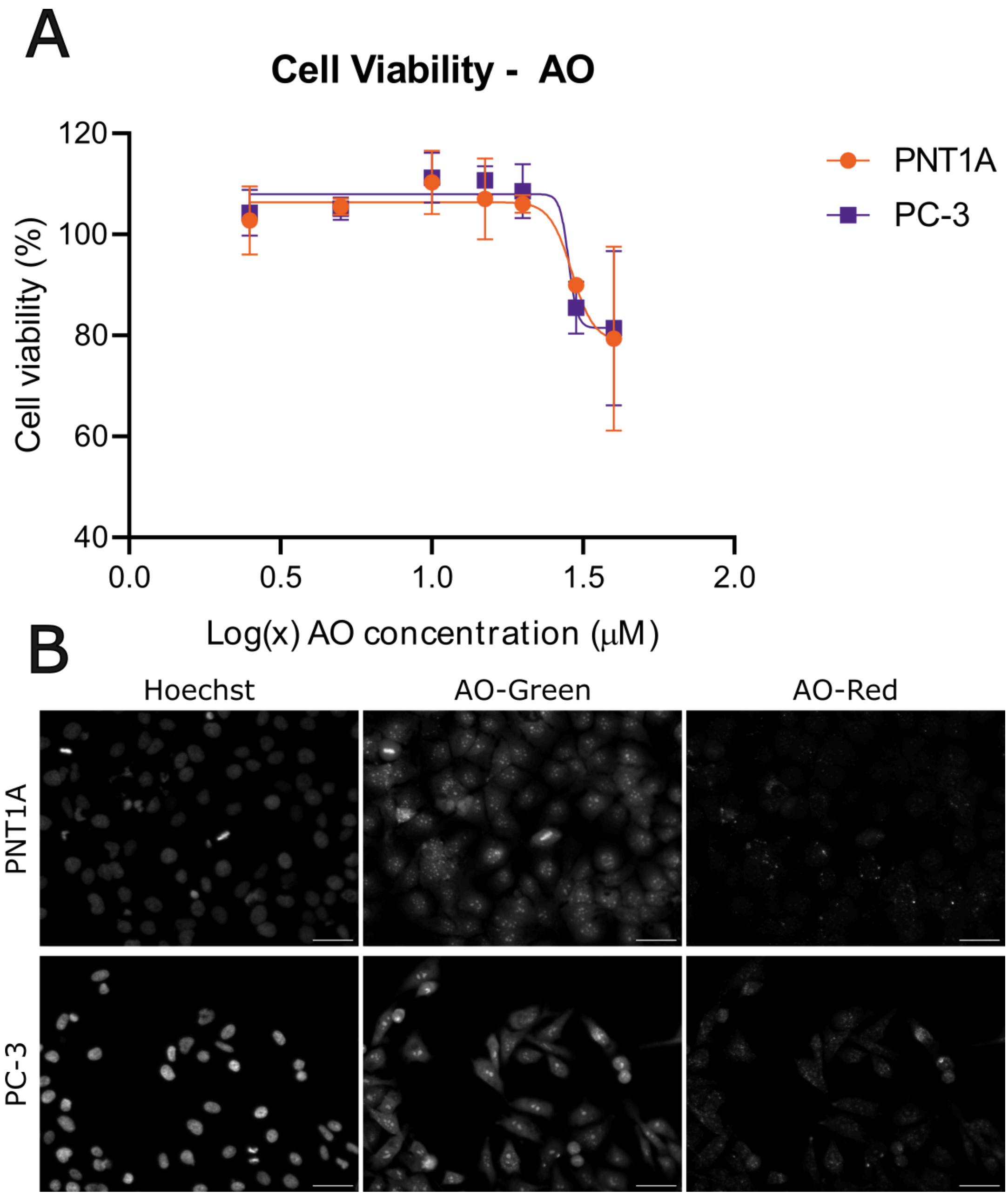


Supplemental Figure 1 - Cell viability of PNT1A and PC-3 cells in the presence of AO assessed by ATP concentration. Cells were treated with AO (2.5, 5.0, 10.0, 20.0, and 40.0 µM) for 4 hours. Then, viability was assessed using CellTiter-Glo® by measuring ATP concentration with luminescence, and images were acquired by Cytation 5, Biotek (20x objective). ATP concentration in the cells was determined by fitting a linear function to the ATP standard curve and calculating the relative viability compared to the negative control. (A) Viability curve for PNT1A and PC3 cells exposed to AO. (B) Representative images for PNT1A and PC-3 cells exposed to 2.5 µM of AO, respectively, represented by the Hoechst,

AO-Green, and AO-Red channels. The lowest concentration tested, 2.5 μM, was selected as it exhibited no cytotoxicity in either cell line. Higher concentrations than 2.5 μM do not decrease cell viability as seen in (A), but cell morphology was visually different than the control. Due to the low concentration of AO, differentiation between the nucleus and cytoplasm was less distinct, cells were stained with Hoechst to facilitate the nuclei segmentation during image analysis. Nonetheless, structures such as the nucleus, nucleoli, vesicles, and cytoplasm were still visible in both cell lines. Optimization of AO concentration should be performed for each cell line. Scale bar = 50 μm. Statistical test performed with two-way ANOVA followed by Sidak's multiple comparisons test: * $p < 0.05$.

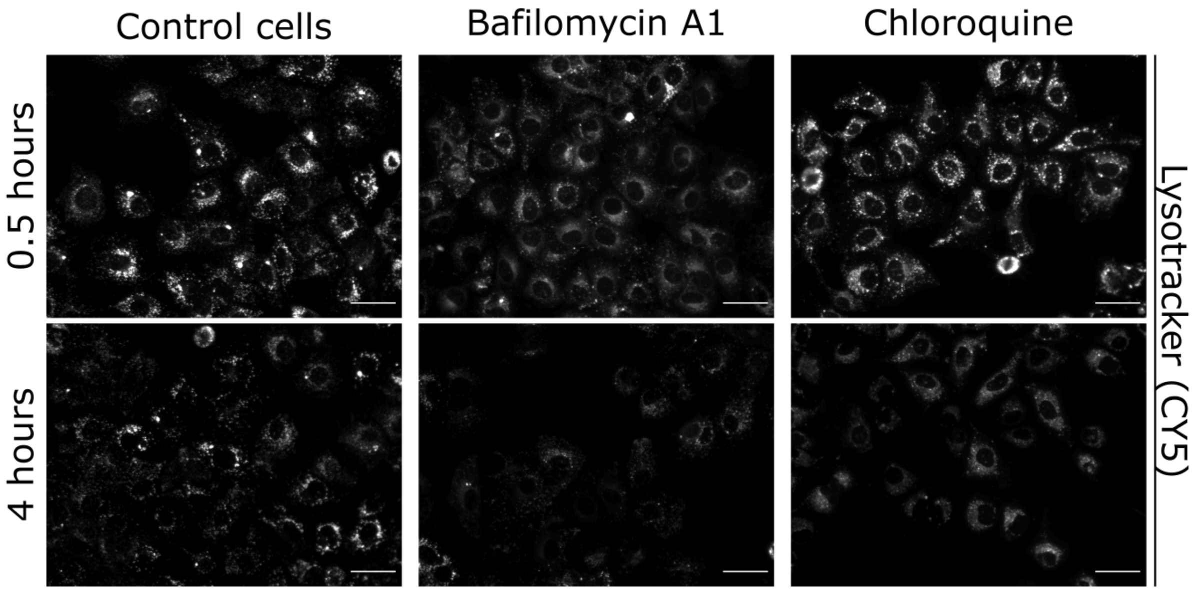


Supplemental Figure 2 - Validation of acidification inhibitors using Lysotracker Deep Red. Huh-7 cells were plated in a 96-well plate at a density of 8000 cells/well. The following day, the cells were treated with solutions containing the inhibitors Bafilomycin A1 1.5 µM or Chloroquine 50 µM added by Lysotracker 75 nM for 0.5 h and 4 h. The images were acquired on the Cytation 5 Hybrid Multidetection Reader (BioTek Instruments, Inc., Winooski, VT, USA using 20× objective. Representative images of the treatments with Lysotracker + inhibitors, acquired in the CY5 channel, represented in grayscale. Scale = 50 µm.

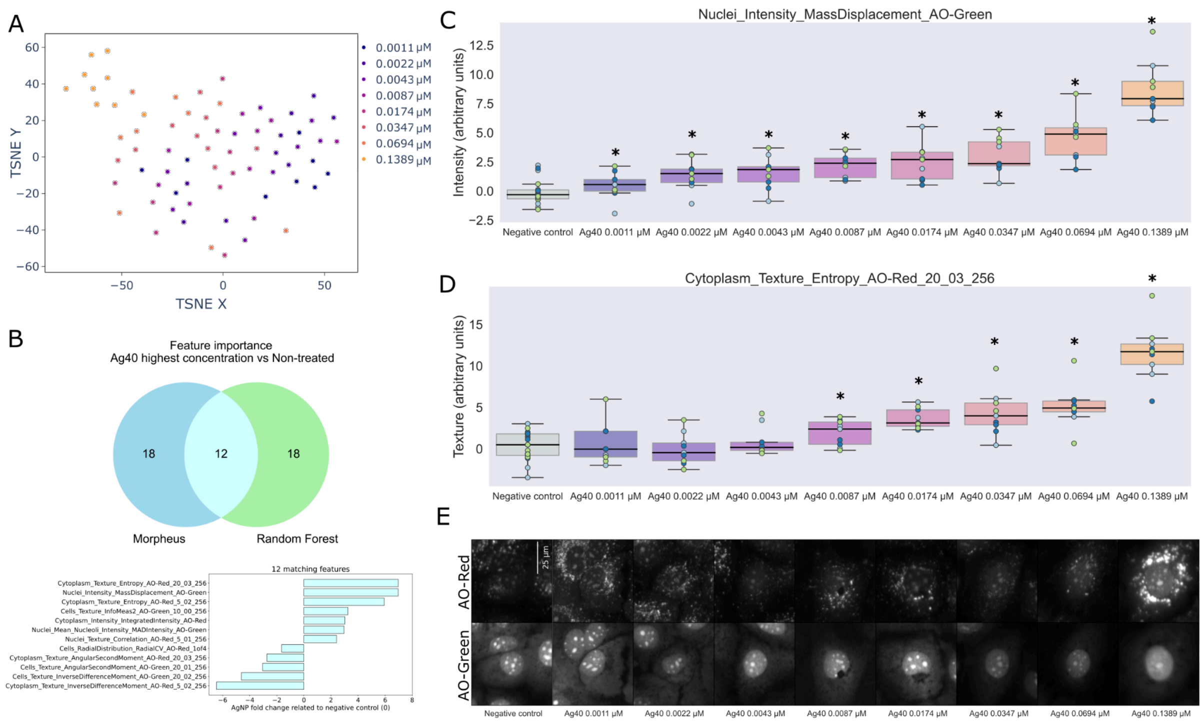


Supplemental Figure 3 - The AO image-based assay detects the dose-response effect on Huh-7 cells induced by silver nanoparticles (AgNP). Huh-7 cells were treated with Ag100 (nanoparticle size of 40 nm) for 24 hours with increasing concentrations of nanoparticles. Then the images were acquired after AO staining (Cytation 5, 20x objective). Objects were segmented using cellpose and features were extracted using CellProfiler; next, features were aggregated by well, normalized, and feature selected using pycytominer, and batch corrected with pyComBat. (A) t-SNE was used for dimensionality reduction and visualization. After 100 iterations, the mean and standard deviation embedding were calculated; (B) Random forest ran for 100 iterations to differentiate between the classes: negative control and Ag40 0.0694 and 0.1389 µM. Then, the mean feature importance and standard deviation were calculated. Morpheus software was used to perform marker selection between the same groups classified in Random Forest. The 30 most important features were selected and compared. Among those, 8 features matched between the methods and were plotted to show the fold change relative to the negative control. The two features highlighted with black boxes are presented in C and D; (C) The intensity of AO-Green in the nuclei; (D) The spatial moment of the nuclei, representing shape, size, rotation, and location of the nuclei object, and (E) Representative images were retrieved from the dataset using k-means clustering (Garcia-Fossa et al., 2023). Experiments were performed in three independent replicates, n =3 and each point in the plots is well-aggregated data, and points are colored by the batch in C and D. Independent t-test calculated using the statannotations library. * p statistical difference < 0.05.

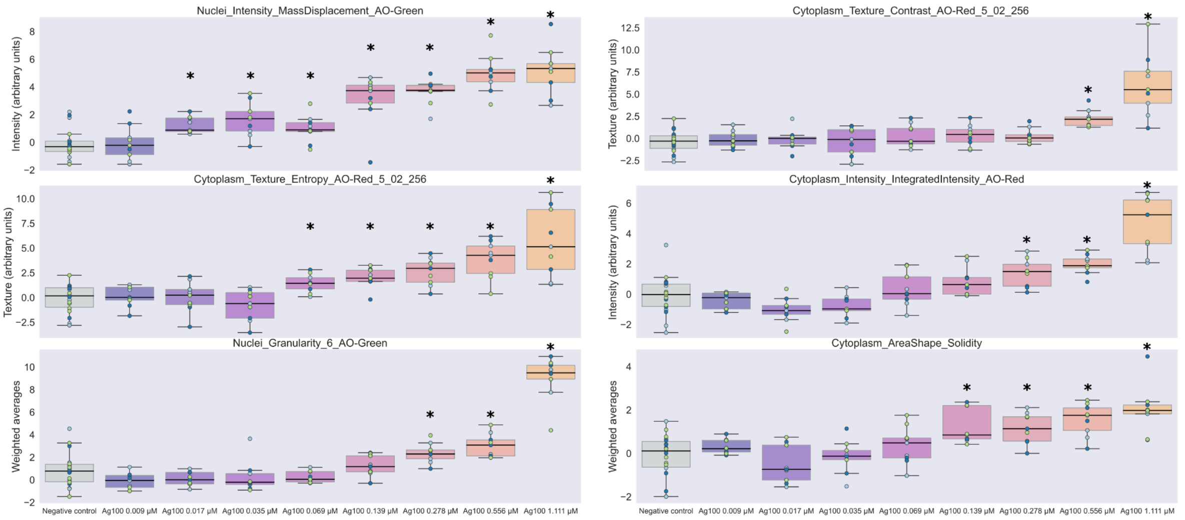


Supplemental Figure 4 - Features that have a dose-response behavior to Ag100 treatment, all showing an increase in overall values relative to the control. The six consistent features identified by both Morpheus and random forest analyses are plotted to illustrate fold changes relative to the negative control. These features include nuclei intensity mass displacement on the AO-Green channel (movement of mass within nuclei); cytoplasm texture contrast on the AO-Red channel (variation in texture within acidic vesicles); cytoplasm texture entropy on the AO-Red channel (randomness in acidic vesicles texture patterns); cytoplasm intensity integrated intensity on the AO-Red channel (overall fluorescence intensity within acidic vesicles in cytoplasm); nuclei granularity on the AO-Green channel (measurement of uniformity and distribution of texture elements); and cytoplasm area shape solidity (compactness of cytoplasm shape).* p statistical difference < 0.05.

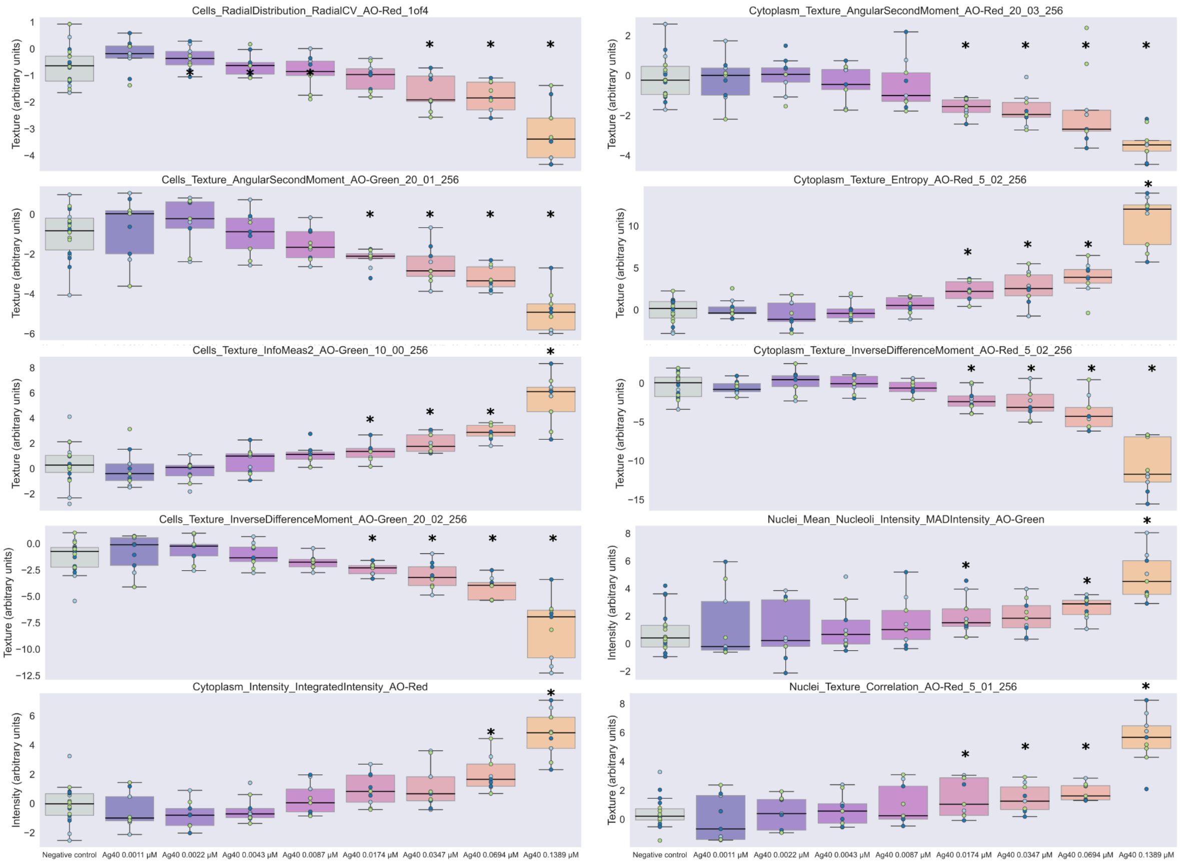


Supplemental Figure 5 - The ten features identified by random forest and Morpheus that have a dose-response behavior to Ag40 treatment are plotted to illustrate fold changes relative to the negative control. These features include: cells radial distribution on the AO-Red channel (distribution pattern of acidic vesicles within the cell); cytoplasm texture angular second moment on the AO-Red channel (local homogeneity of acid vesicles in cytoplasm texture in angular patterns); cell texture angular second moment on the AO-Green channel (local homogeneity of cell texture in angular patterns); cytoplasm texture entropy on the AO-Red channel (randomness in cytoplasmatic acidic vesicles texture patterns); cells texture infomeas2 on the AO-Green channel (measure of texture complexity in cells); cytoplasm texture inverse difference moment on the AO-Red channel (local homogeneity of cytoplasmatic acidic vesicles texture in pixel intensity differences); cells texture inverse difference moment on the AO-Red channel (local homogeneity of acidic vesicles texture in pixel intensity differences); nuclei mean nucleoli intensity mad intensity on the AO-Green channel (average intensity of nucleoli within nuclei); cytoplasm intensity integrated intensity on the AO-Red channel (overall acidic vesicles fluorescence intensity within the cytoplasm); nuclei texture correlation on the AO-Red channel (correlation between texture elements within nuclei).* p statistical difference $< 0.05$.

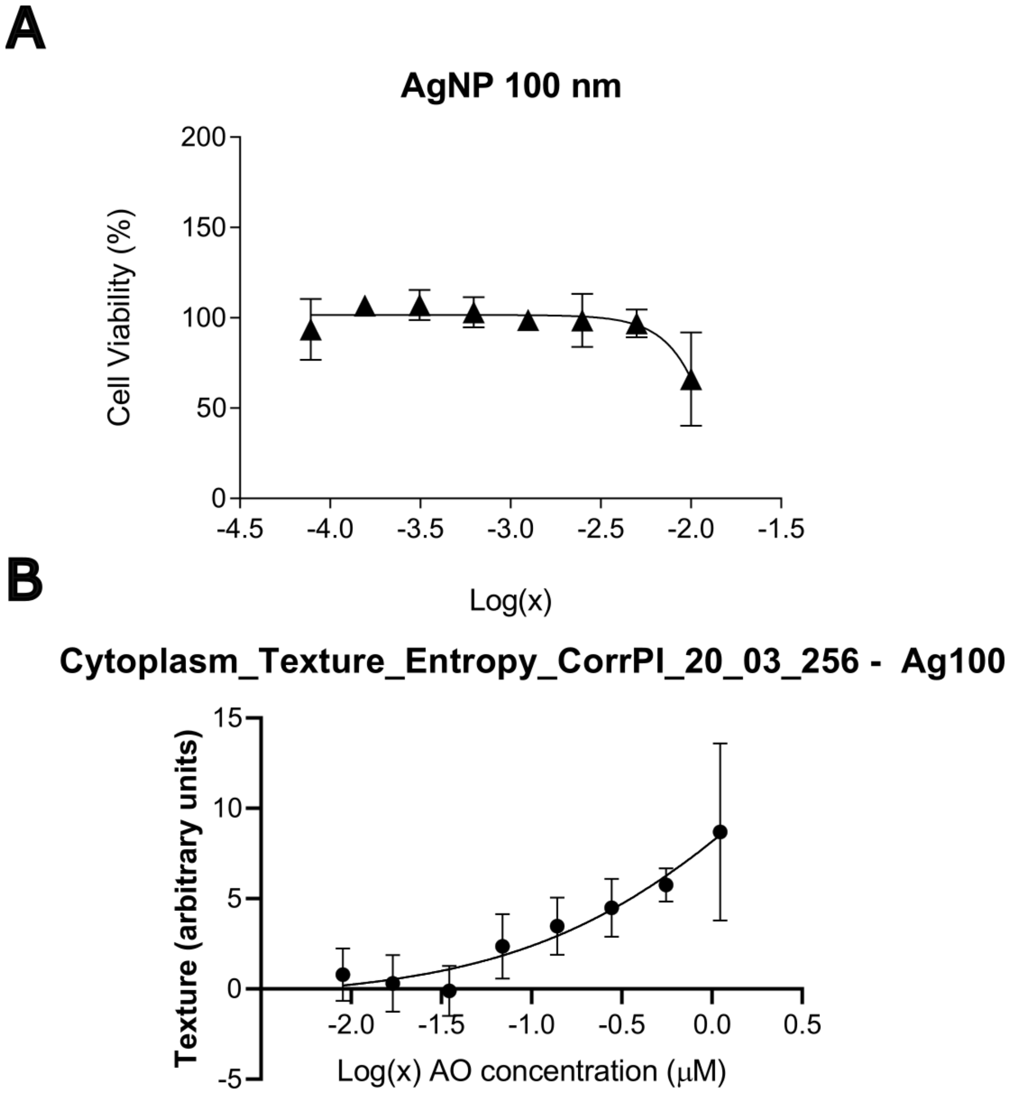


Supplemental Figure 6 - Comparison of IC50 calculation using MTT data or feature data. (A) The cells were treated with enhancing concentrations of AgNP 100 nm for 24 hours. Then, MTT 5 mg/mL was added and incubated for 2 hours. MTT absorbance was read on 570 nm using Cytation 5 Biotek. (B) The feature median values for each AgNP concentration were used to calculate the IC50.

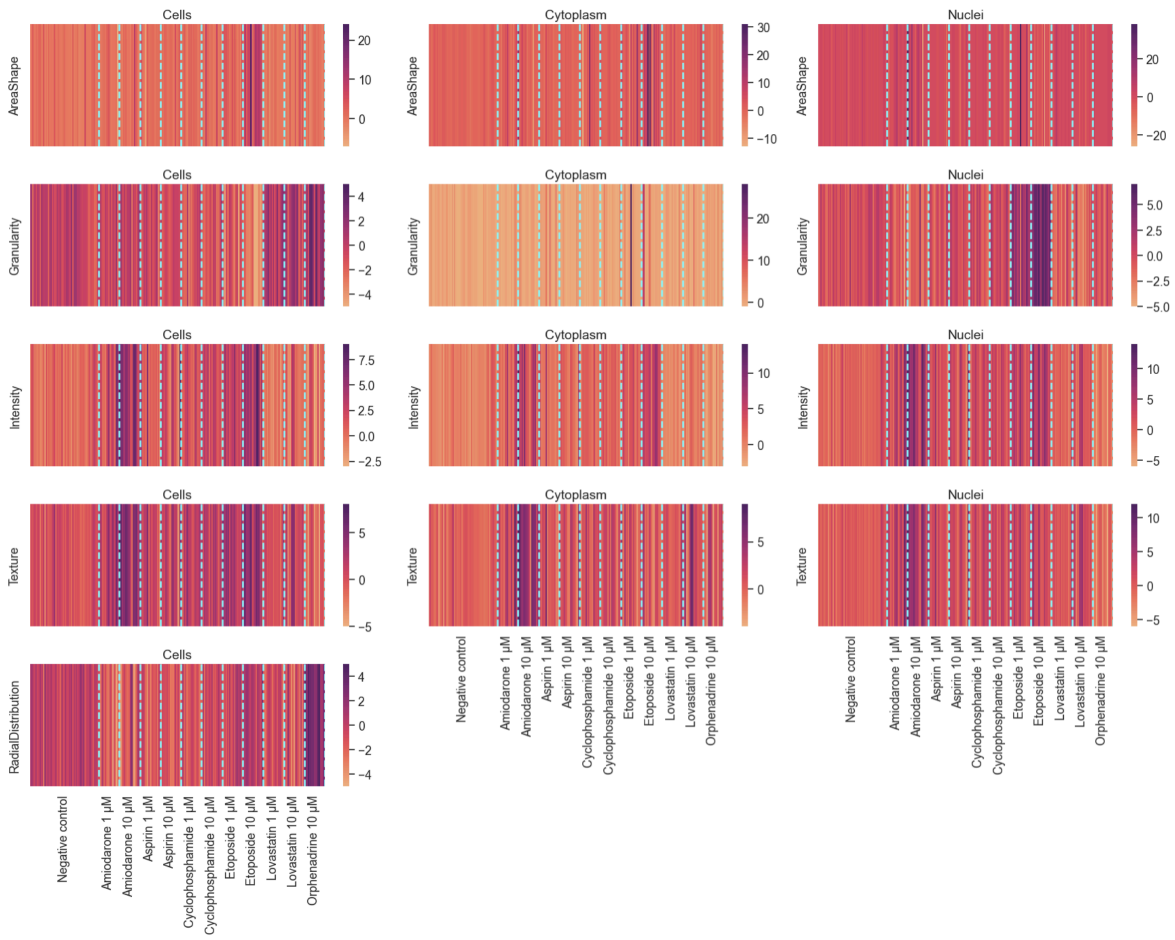


Supplemental Figure 7 - Features extracted from DILI-drugs assay after feature selection and split by categories and compartments. For the features in each category and each compartment, we calculated the principal component 1 to be a summary of all the features in that category. All features were normalized to the negative control, which has zero value in this matrix. The treatments induce a variation for each category that can be lower or higher than the negative control.

A

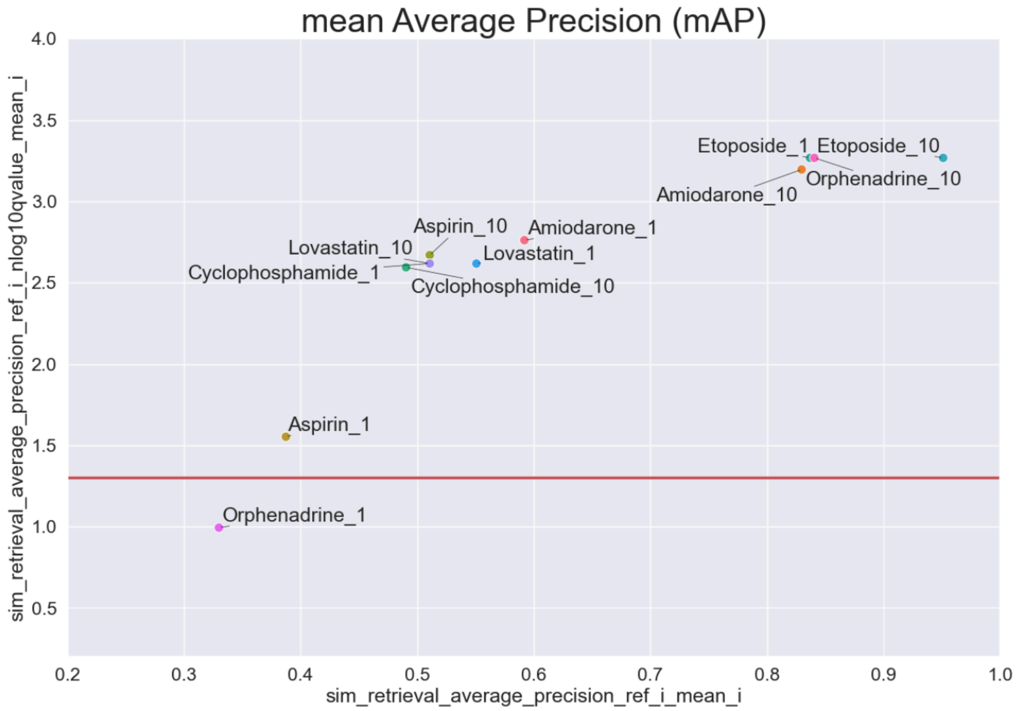


B

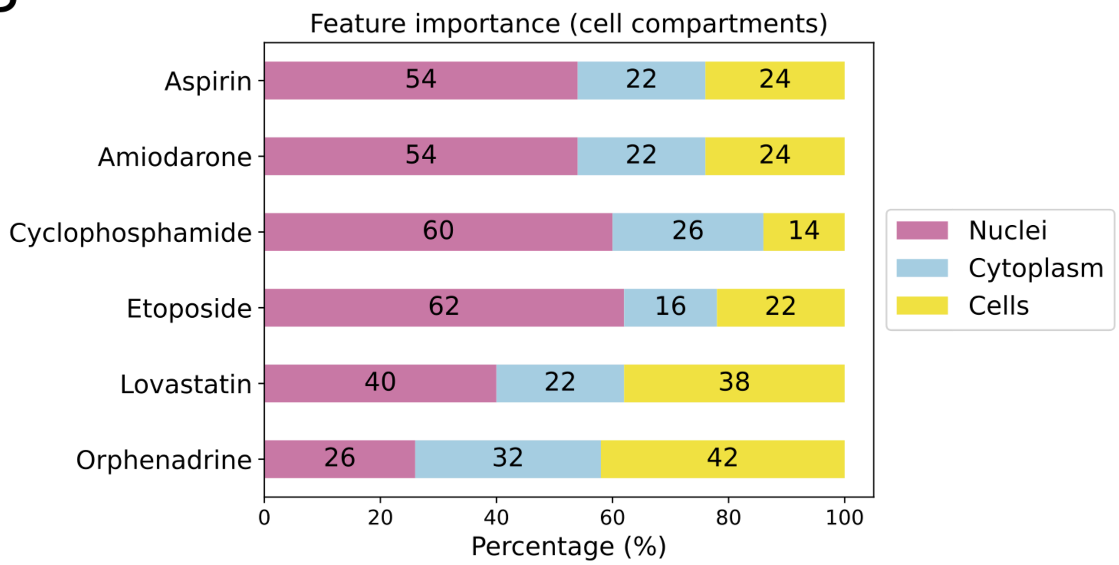


Supplemental Figure 8 - Evaluation of technical replicability of the profiles and compartments' contribution for the 30 most important features. (A) Retrievable phenotypes are above the threshold (red line in x-axis, significance threshold 0.05). These compounds' ability to retrieve themselves relative to the negative control is defined by their mean Average Precision (mAP) in the x-axis, and the q-value in the y-axis helps define the threshold of 0.05; (B) The contribution of each compartment for the most important features: Nuclei, Cytoplasm, and Cells.

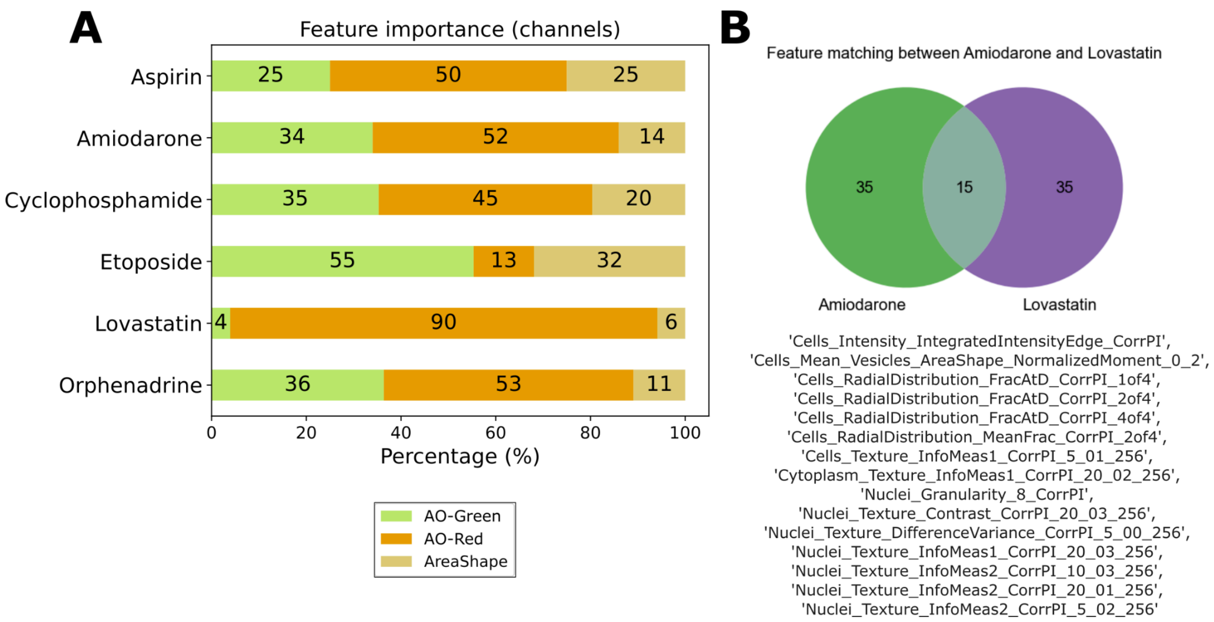


Supplemental Figure 9 - Feature matching between amiodarone and lovastatin. After selecting the 50 most significant features using Random Forest and Morpheus software, the matching features between the compounds were obtained. (A) The percentage of the 50 most important features that belong to AO-Green, AO-Red, or AreaShape channels. For lovastatin and amiodarone, the percentage of features associated with the AO-Red is high. (B) All matching features are related to the AO-Red channel except for one associated with the shape of the vesicles.

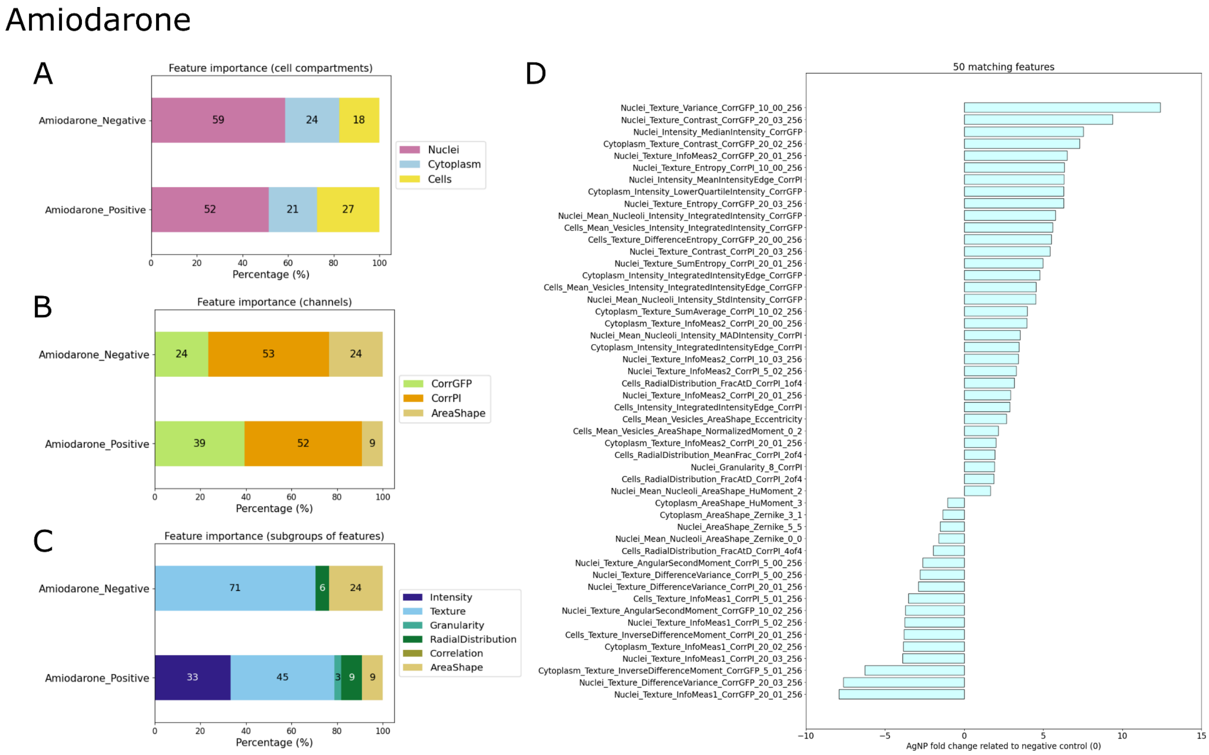


Supplemental Figure 10 - Key features defining amiodarone clustering. After selecting the 50 most significant features using Random Forest and Morpheus software, the median value of each feature was subtracted from the median value of the negative control. Features were then categorized as either negative or positive relative to the negative control. The results are presented as follows: (A) cell compartments, (B) channels, (C) subgroups of features, and (D) the complete list of features along with their fold changes relative to the negative control.

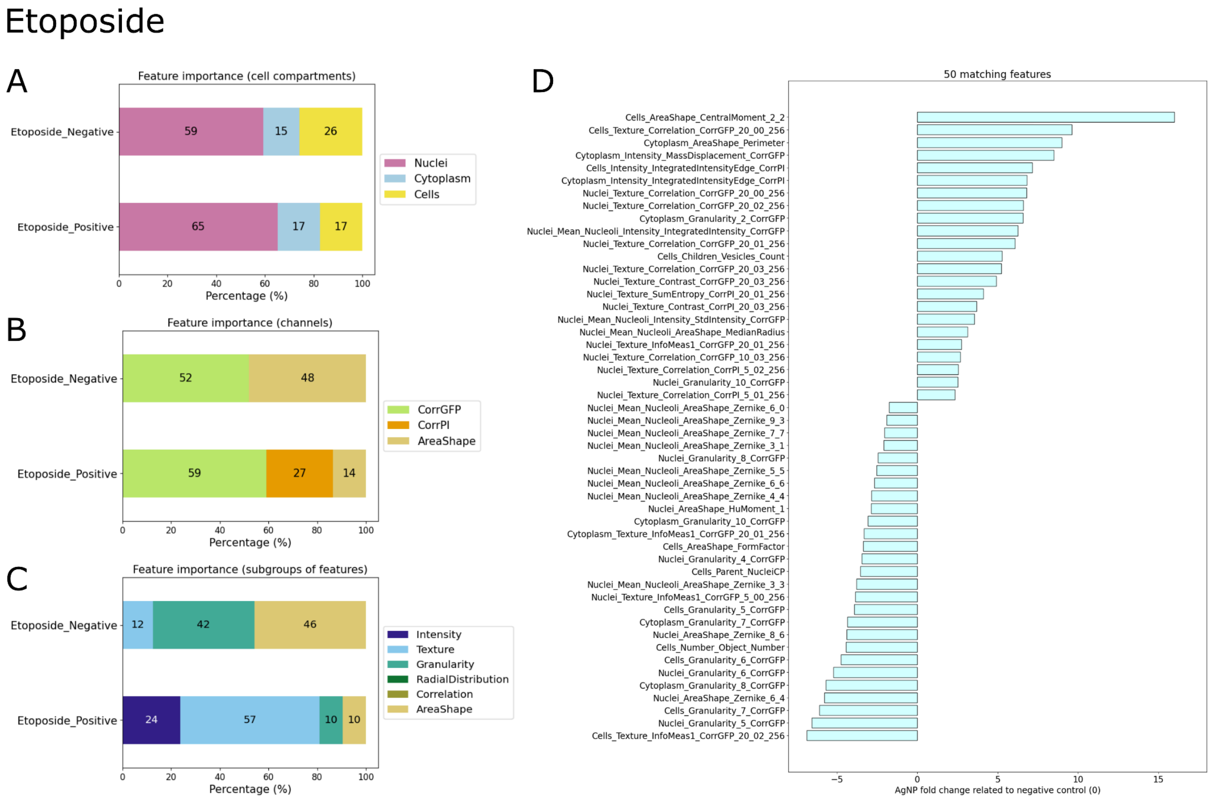


Supplemental Figure 11 - Key features defining etoposide clustering. After selecting the 50 most significant features using Random Forest and Morpheus software, the median value of each feature was subtracted from the median value of the negative control. Features were then categorized as either negative or positive relative to the negative control. The results are presented as follows: (A) cell compartments, (B) channels, (C) subgroups of features, and (D) the complete list of features along with their fold changes relative to the negative control.

### Orphenadrine

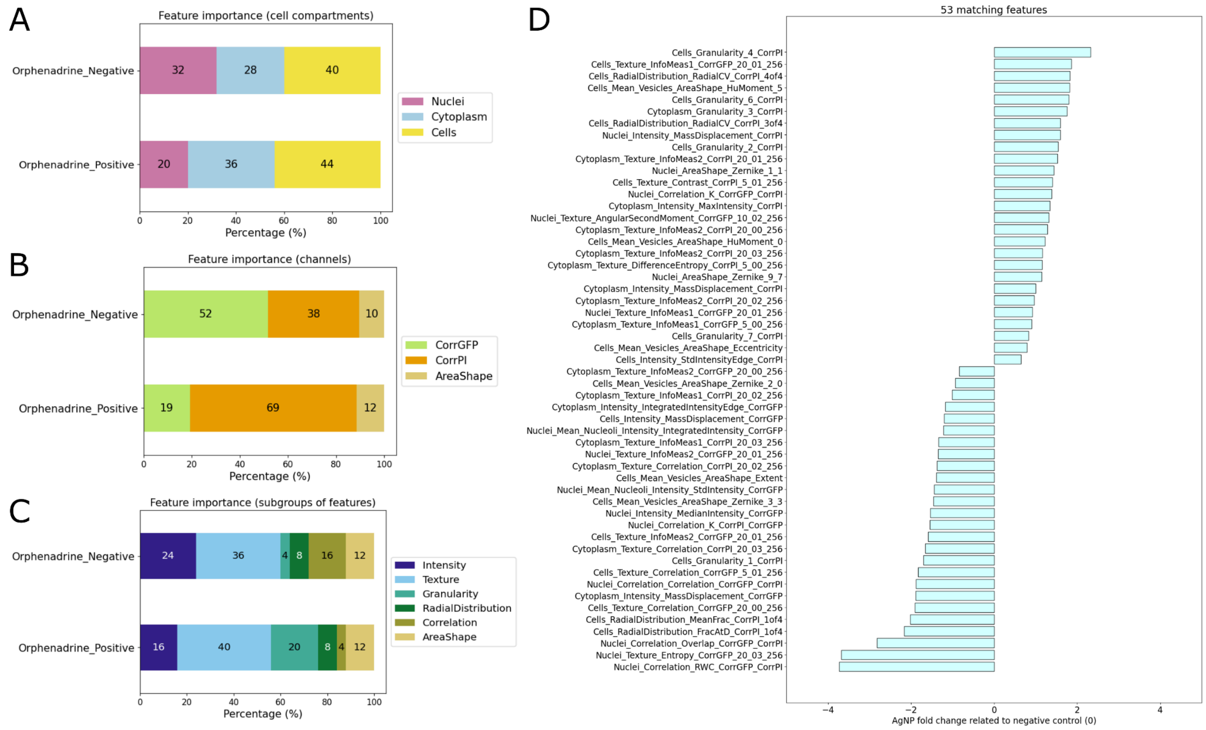


Supplemental Figure 12 - Key features defining orphenadrine clustering. After selecting the 50 most significant features using Random Forest and Morpheus software, the median value of each feature was subtracted from the median value of the negative control. Features were then categorized as either negative or positive relative to the negative control. The results are presented as follows: (A) cell compartments, (B) channels, (C) subgroups of features, and (D) the complete list of features along with their fold changes relative to the negative control.
